# Deeper insights into long-term survival heterogeneity of Pancreatic Ductal Adenocarcinoma (PDAC) patients using integrative individual- and group-level transcriptome network analyses

**DOI:** 10.1101/2020.06.01.116194

**Authors:** Archana Bhardwaj, Claire Josse, Daniel Van Daele, Christophe Poulet, Marcela Chavez, Ingrid Struman, Kristel Van Steen

## Abstract

**Background:** Pancreatic ductal adenocarcinoma (PDAC) is categorized as the seventh leading cause of cancer mortality worldwide. Its predictive markers for long-term survival are not well known. Therefore, it is interesting to delineate individual-specific perturbed genes when comparing long-term (LT) and short-term (ST) PDAC survivors, and to exploit the integrative individual- and group-based transcriptome profiling.

**Method:** Using a discovery cohort of 19 PDAC patients from CHU-Liège (Belgium), we first performed differential gene expression (DGE) analysis comparing LT to ST survivor. Second, we adopted unsupervised systems biology approaches to obtain gene modules linked to clinical features. Third, we created individual-specific perturbation profiles and identified key regulators across the LT patients. Furthermore, we applied two gene prioritization approaches: random walk-based Degree-Aware disease gene prioritizing (DADA) method to develop PDAC disease modules; Network-based Integration of Multi-omics Data (NetICS) to integrate group-based and individual-specific perturbed genes in relation to PDAC LT survival.

**Findings:** We identified 173 differentially expressed genes (DEGs) in ST and LT survivors and five modules (including 38 DEGs) showing associations to clinical traits such as tumor size and chemotherapy. DGE analysis identified differences in genes involved in metabolic and cell cycle activity. Validation of DEGs in the molecular lab suggested a role of REG4 and TSPAN8 in PDAC survival. Individual-specific omics changes across LT survivors revealed biological signatures such as focal adhesion and extracellular matrix receptors, implying a potential role in molecular-level heterogeneity of LT PDAC survivors. Via NetICS and DADA we not only identified various known oncogenes such as CUL1, SCF62, EGF, FOSL1, MMP9, and TGFB1, but also highlighted novel genes (TAC1, KCNH7, IRS4, DKK4).

**Interpretation:** Our proposed analytic workflow shows the advantages of combining clinical and omics data as well as individual- and group-level transcriptome profiling. It suggested novel potential transcriptome marks of LT survival heterogeneity in PDAC.

**Funding:** Télévie-FRS-FNRS

## Introduction

Pancreatic ductal adenocarcinoma (PDAC) accounts for 90% of pancreatic tumors.^1^ It is the 4th leading cause of cancer-related death worldwide, while remaining the most lethal among digestive cancers.^2^ PDAC has a complex and dense tumor microenvironment that poses a significant barrier to treatment administration.^3^ Various factors shape the outcome for complex diseases leading to perturbations of a complex intracellular network.^4^ Disease-relevant genes typically do not operate on their own but may be connected to each other and known disease associated genes of interest.^5^ Network approaches that allow integration with regulatory factors are required to fully map complex diseases, including PDAC.

For PDAC, the overall survival (OS) of patients may be coupled to the mutational status of *KRAS* (Kirsten rat sarcoma viral oncogene) as well as several morphological features.^6^ Also, multiple miRNAs and transcription factors influence metastasis and OS time of PDAC patients.^7, 8^ Due to the high lethality of PDAC, intensive research is needed to unravel roots of causes for PDAC survival in general and long-term (LT) versus short-term (ST) survival in particular. In the literature, several criteria for LT and ST survival exist : ST (resp. LT) as surviving ≤ 8 (resp. ≥ 8 months)^9^; LT survival as ≥ 10 years^10^; ST (<14 months) and very long-term (≥ 10 years) of survival.^11^ Very little information is available about regulatory mechanisms involved in the context of <12 months and ≥36 months of survival within European populations. We aim to fill this gap and to explore PDAC survival mechanisms by making use of genomics data and by integrating a variety of gene prioritization methods.

Multiple questions are of interest, including ‘How do LT and ST PDAC survivors differ from each other’ and ‘Which survival group is most heterogeneous in terms of transcriptome signatures’. PDAC is featured with intra-tumoral heterogeneity.^12^ In general, heterogeneity poses a significant challenge to personalized treatments for PDAC.^13^ Previous classification studies paved the path to a better classification of patients with PDAC based on molecular pathology information^14^, molecular features^15^ and defined five PDAC subtypes, showing associations with patient outcomes.^16^ The identification of subgroups by looking into a perturbed profile of each individual might be another interesting approach. Typically, such (molecular) subtyping analyses require relatively large sample sizes. Alternative and more elaborate approaches are required, better exploiting and combining individual-level and group level profiling, to address the aforementioned questions.

Pathological findings with tumor cells suggest an abundance of gene regulatory networks (GRNs) in humans for various cancers including, breast^17, 18^, prostate^19^, and PDAC cancer.^20^ Network biology approaches have the potential to identify key regulators that are responsible for molecular heterogeneity giving rise to LT and ST PDAC survivor subgroups. Weighted Gene Co-Expression Network Analysis (WGCNA) is such an approach and enables the identification of gene modules and their associations with clinical measurements^21^. For the identification of PDAC key regulators, more work is needed to exploit gene connectivity with earlier identified disease genes via the use of protein interaction networks (PPIs).

The current study teases out PDAC survival associated genes, with a focus on LT survivors (≥ 36 months survival; in contrast to ST survival defined as ≤ 12 months survival) and individual-to-individual differences in whole transcriptome profiles. To this end, we introduced and implemented a flexible and interpretable omics integrative analysis framework, involving a series of group-level and individual-level viewpoints.

## Methods

The study’s analytic workflow is depicted in Figure 1 and described in more detail in appendix pp 2-5.

**Figure 1:**
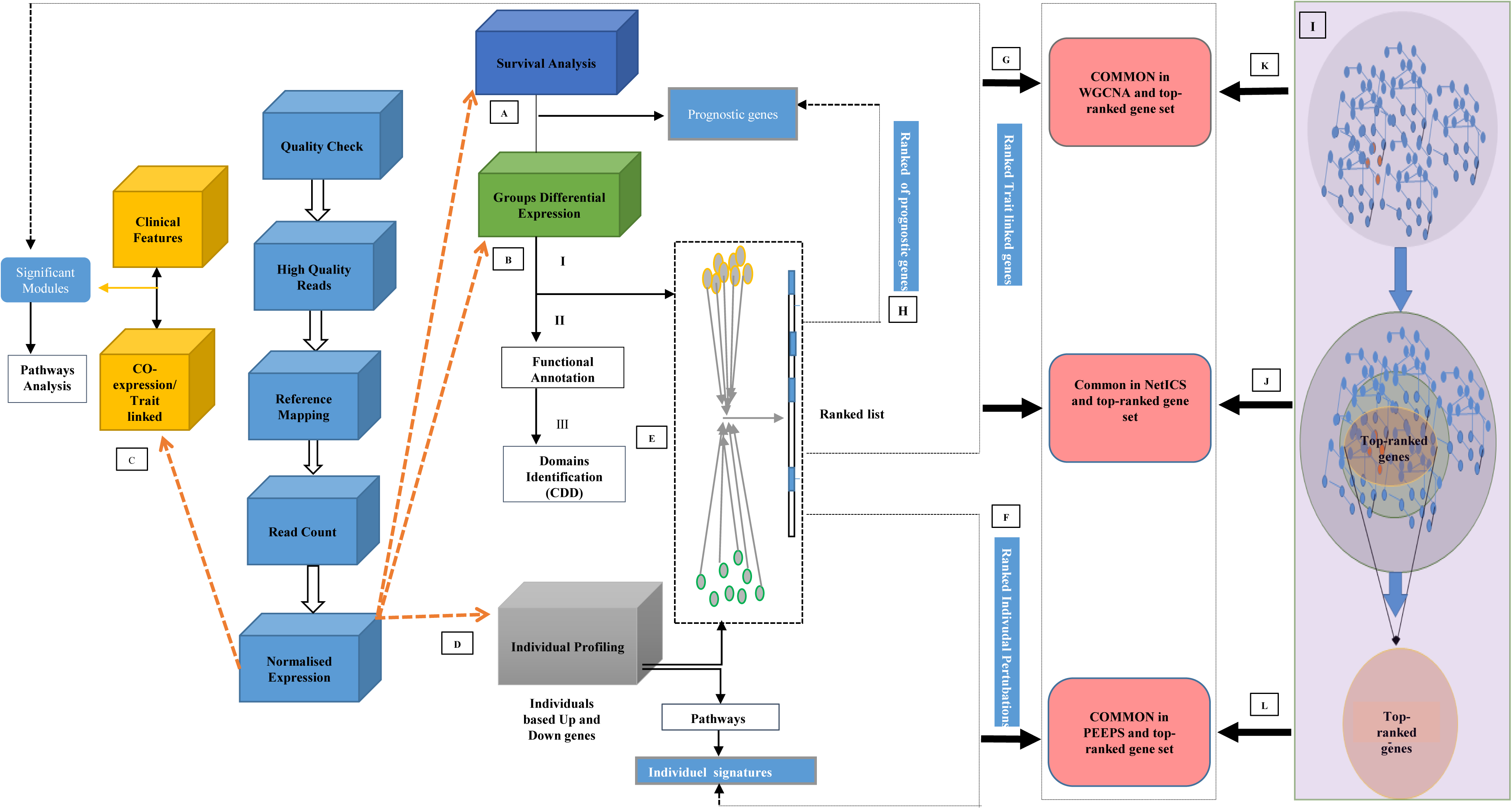
Flexible and interpretable omics integrative framework for RNA-seq data collected on two groups of patients, exemplified on PDAC ST/LT survival. RNA-seq quality-controlled data are inputted for A) Survival analysis; B) Group-based differential analysis via DESeq2^60^; C) Weighted gene co-expression network analysis WGCNA^21^; D) Individual-based differential analysis (appendix pp 2-5); E) Genes are ranked based on the integration of individual and group-based differentially expressed genes via NetICS^61^; F-H) NetICS specific top 1% ranked genes are traced back in multiple previous analyses (A through E); I) DADA^5^ analysis starting from disease genes; J-L) DADA specific top 1% ranked genes are traced back in previous analyses (A through E).

## Results

### Patients characteristics

All patients were divided into ST (≤12 months) and LT (≥36 months) survival groups (resp. STS and LTS), as summarized in Figure 2A. A total of 19 patients, comprising 10 STS and 9 LTS, met our inclusion criteria (appendix pp 2;9-11).

**Figure 2:**
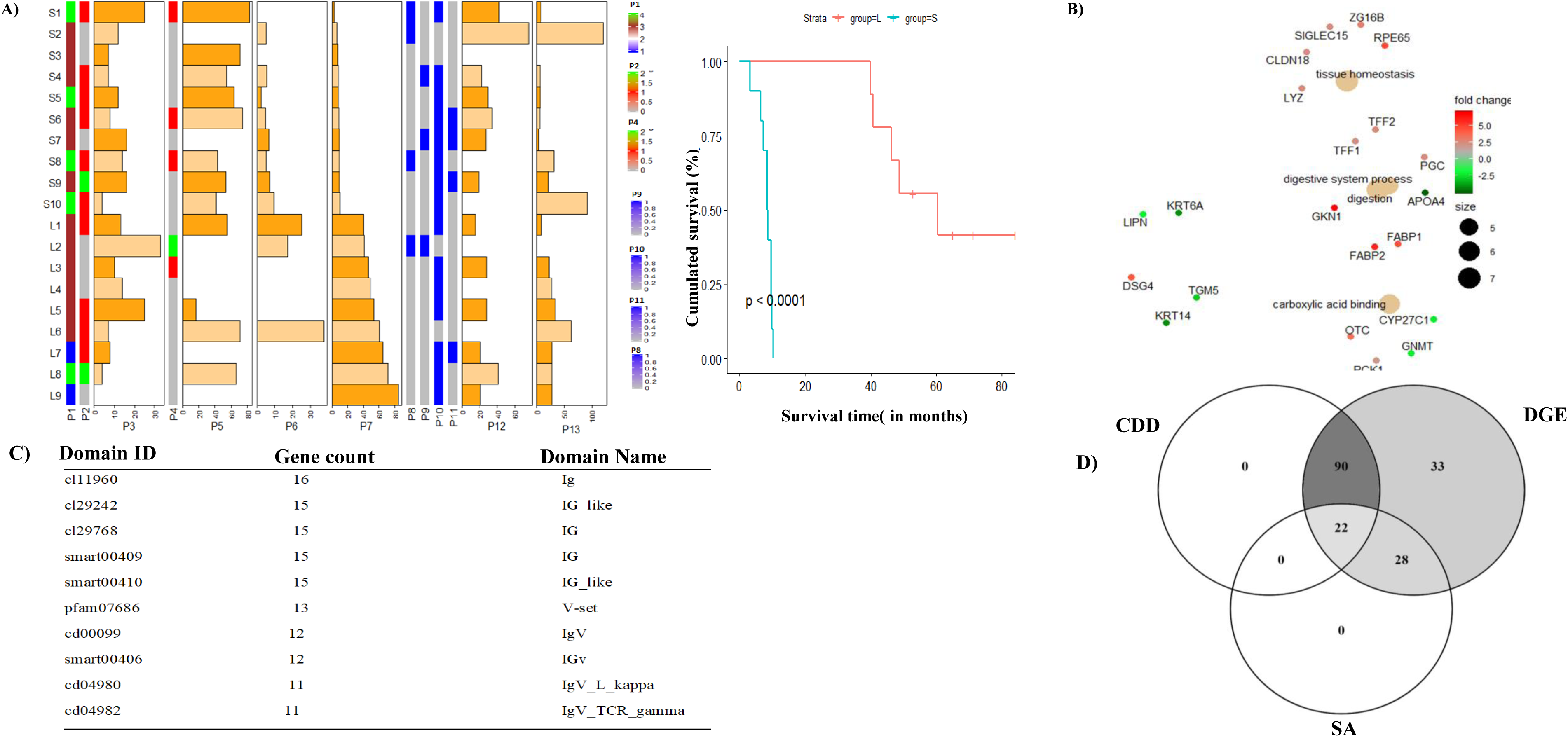
Overall Kaplan–Meier survival analysis of the ST and LT PDAC cohorts: A) Patient characteristic data for a selection of PDAC relevant traits are shown as mixed bar and heat map plot. P1 to P13 refer to patient specific clinical traits analyzed in this study (selective data has been shown in plot; full details given in appendix pp 9-11). P1 indicates Tumor stage (from 1 to 4). P3, P5, P6, P7, P12 and P13 indicate the frequency of number of nodes analyzed, time between surgery and chemotherapy (in days), disease free survival, OS (in months), tumor size by imagery (in mm) and Time between imagery and surgery, respectively. Remaining P2, P4, P8, P9, P10, P11, refers to status of N stage, surgical margin invaded by tumor cells, vascular resection, re-hospitalization after surgery, vascular contact, and artery contact, respectively. Here 0 and 1 indicate no and yes, reps. P7 clinical trait denotes overall survival and was used for the development of the Kaplan-Meier survival curves for short-term (ST) and long-term (LT) PDAC Survivors (STS: S1 to S10; LTS: L1 to L9); B) Identification of significant gene ontology of associated up and down-regulated DEGs and their relevant functions. Up and down-regulated genes are highlighted with red and green dots, respectively. The size of data points increases with increased significance (uncorrected for multiple testing – see appendix pp 3); C) Top-ranked conserved domains in differentially expressed gene sets; D) Venn-diagram showing the number of identified genes that are common to or different in multiple first-line analysis strategies (CDD: conserved domain database analysis, DGE: differential gene expression analysis, SA: survival analysis (appendix pp 3)).

### Differential gene expression analysis and functional follow-up

RNA was extracted from FFPE tissues and a quality check was performed for paired-end sequencing. The long non-coding gene *MIR205HG* was the topmost differentially down-regulated in the LTS group (p-value=0.008), while the protein coding gene *GKN1,* which encodes for gastrokine1, was the topmost differential up-regulated in LTS (p-value=1.25E-05). The gene ontology analysis linked the genes with the highest expression levels in LTS to the digestive system and lowest expression levels in LTS to the carboxylic binding activity (Figure 2B). Specific domain structures of genes play a significant role in gene regulation and expression. The conserved domain analysis resulted in 112 genes containing at least one domain (Figure 2C; appendix pp 3). Sixteen genes contained an Ig domain, followed by a V-set domain. Later, a unique set of domains was identified in both up- (IgC, IG_like, Trypsin) and down-regulated genes (F-Box, IRS, PKc_MAPKK, IgV) suggesting these genes’ regulatory role in PDAC survival mechanisms. Fifty survival genes were identified from all DEGs. We observed 22 DEG genes containing at least one domain that overlapped with the survival gene set (Figure 2D). RT-qPCR confirmed the differential expression observed in LTS versus STS for the genes represented in appendix pp 33. Among them, DEGs *REG4* and *TSPAN8* were validated in lab via RT-qPCR analysis (appendix pp 5:section 3.1).

### Group-level survival heterogeneity: Significant clinically relevant modules and their corresponding 3D architectures

All 19 samples with clinical information and gene expression were included in WGCNA (appendix pp 3). Genes with similar expression were grouped into gene modules via average linkage hierarchal clustering. By use of a dynamic tree-cutting algorithm, a total of 96 distinct co-expression modules were identified. Correlated modules were merged with a cut-off height of 0.25, resulting in 35 modules containing 66 to 2010 genes per module. Module M34 was the smallest module consisting of 66 genes, whereas M8 was the largest module comprising 2010 genes (Figure 1:C; Figure 3A). The identified 35 modules covered 97 percent of the 18880 input genes. For those 35 modules, we derived the corresponding module eigengenes (MEs).

**Figure 3:**
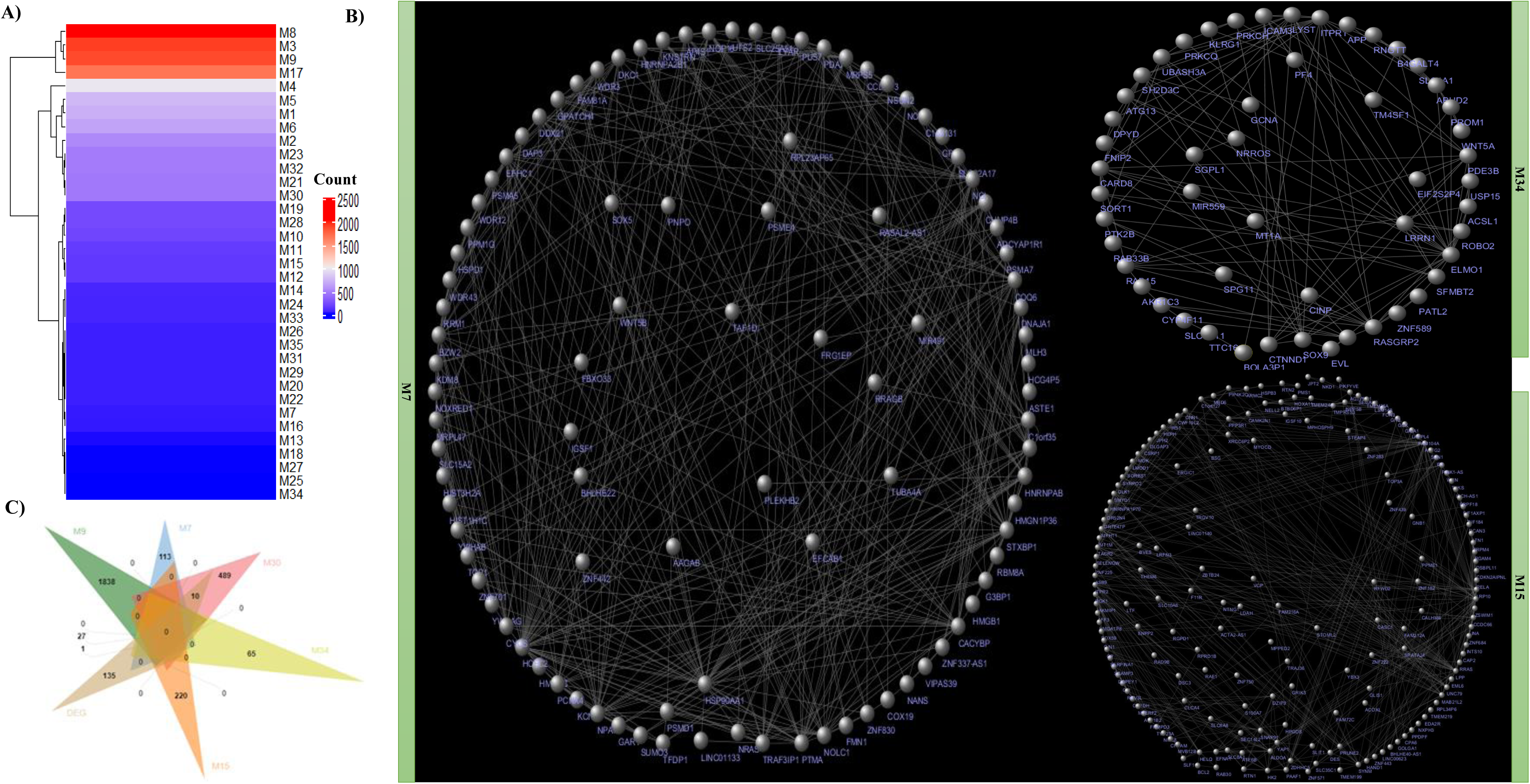
Clinical relevance of gene co-expression modules: A) Heatmap indicating the number of genes involved in each WGCNA-derived gene module; B) Network topology of three modules (M7, M15, M34), where nodes are genes and connections among nodes represent gene-gene interactions. In each network, the gene names are indicated in the circular layout as derived from Cytoscape.^62^; C) Venn diagram indicating the common genes between the identified significant DEGs and the five previously identified clinically relevant modules.

Association of clinical features with dysregulated genes may help to clarify genes important for disease development. All identified DEGs (173 in total) were distributed in 25 modules. Five modules had a significant correlation with clinical phenotypes (adjusted p-value <0.05): M7, M9, M15, M30, and M34 (appendix pp 34). Module M9 was found to be significantly associated with tumor size (r^2^=0.72, adjusted p-value=0.01) and T stage (r^2^ = 0.68, adjusted p-value=0.03). M9 consisted of the highest number of DEGs (27 genes). Two other modules, M7 (r^2^=0.73, adjusted p-value=0.01) and M30 (r^2^=0.71, adjusted p-value=0.02), were negatively associated with time between surgery and chemotherapy clinical traits. M30 contained 10 DEGs. Module M34 was significantly associated with tumor size by imagery (r^2^=0.67, adjusted p-value<0.05). Interestingly, two modules were significantly associated with chemotherapy: a positive association for M15 (r^2^=0.68, adjusted p-value=0.04) and a negative association for M9 (r^2^= −0.68, adjusted p-value<0.04). Gene-gene interactome networks were developed for clinically relevant modules M7, M15 and M34 (Figure 3B). The overlap between DEGs and genes in five modules (M7, M9, M15, M30, M34) is shown in a Venn-Diagram (Figure 3C), and identified 27, 10, and 1 gene as part of M9, M30, and M34, respectively.

### Group-level survival heterogeneity: Functional analysis of clinically relevant modules

Clinically pertinent gene modules were functionally analyzed in Cytoscape with the ClueGO plug-in (appendix pp 3-4) to visualize functionally grouped network. Module M9 was linked to 33 significant pathways (adjusted p-value <0.05) distributed over ten groups, such as extracellular matrix (ECM) organization (86 genes) and collagen formation (37 genes) (Figure 4A). Genes regulating the cell cycle and modulating ECM at molecular or cellular levels have been linked to cancer drug targeting and cancer cell plasticity.^32^ M7, also negatively associated with chemotherapy, contained 91 significant pathways, distributed into three groups, such as proteasome (4 genes) and the regulation of *RAS* by *GAP*s (5 genes) (Figure 4B). Module M15, positively associated with ‘chemotherapy’, was enriched with 11 significant pathways distributed into five groups (Figure 4C). More detail is given in appendix (pp 5: section 3.2)

**Figure 4:**
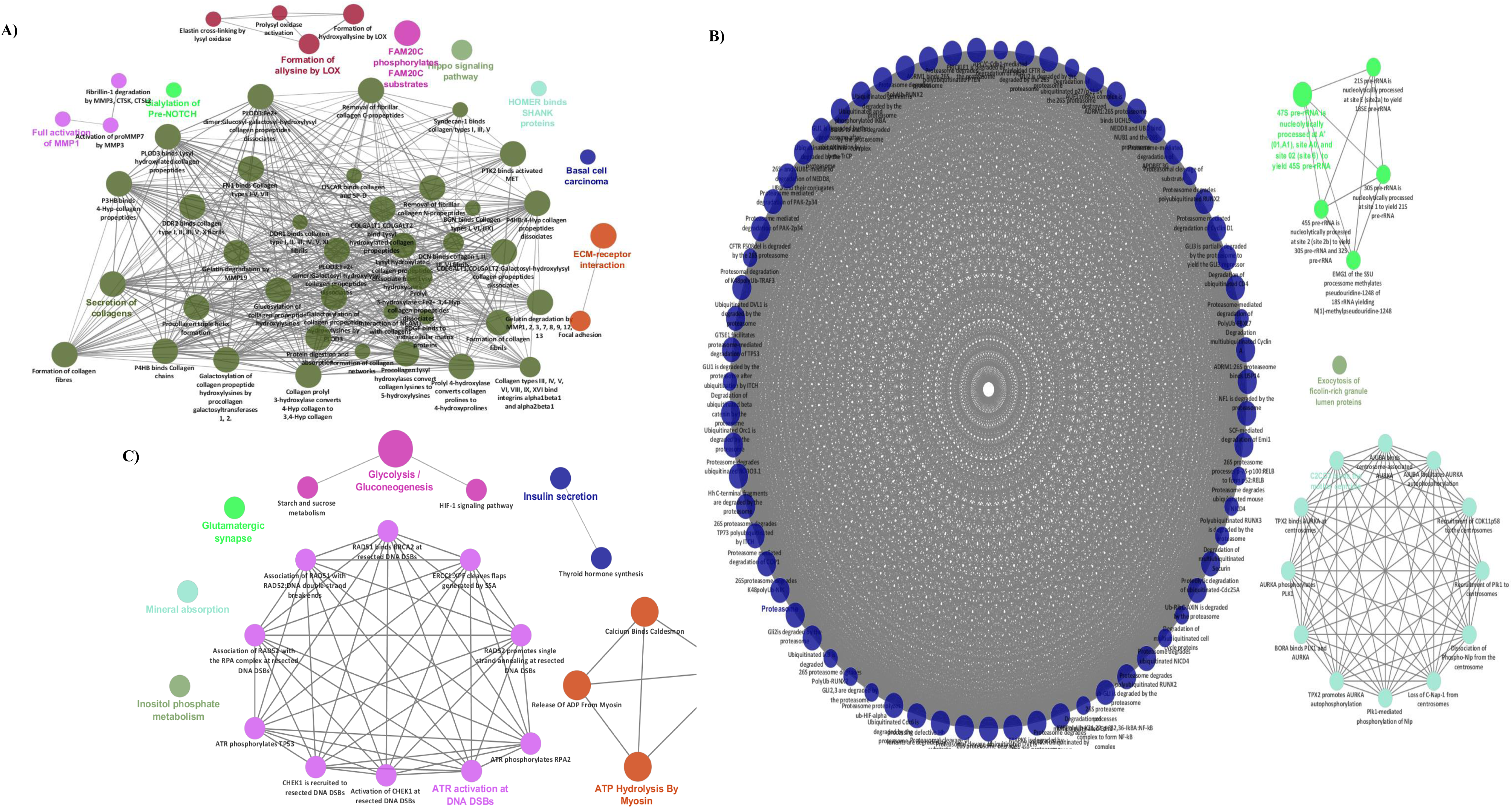
Functional follow-up of clinically relevant gene expression modules: A) Ten groups for module M9 comprising 33 significantly linked pathways; B) Three groups identified in the M7 modules; C) Depiction of the five groups identified in M15; For A-C, redundant groups with >50% overlap were merged. Each node in the network represents an enriched term; the size of each node follows the extent of enrichment significance. Connection among different nodes are based on kappa scores (≥0.4), as available from ClueGO.

### Individual-specific survival heterogeneity: Quantification of heterogeneity between individual transcriptome profiles

To assess heterogeneity in LT survival patients, we constructed individual perturbation expression profiles (PEEPs)^22^ (appendix pp 4). It resulted in 6336 significantly perturbed genes across LT PDAC survivors (Figure 1:D; Figure 5A). The frequency of disrupted genes in each LT survivor Li (i= 1,…,9) was L1:12, L2:1412, L3:43, L4:474, L5:179, L6:319, L7:957, L8:150 and L9:2789 (Figure 5A). Various genes were uniquely perturbed in one LTS patient only. Only one group-wise DEG was shared among 3 LT survival subjects, namely TNNI3. Also, at most six DEGs (IRS4, KLRC3, CLDN18, NPY, CNTN6, TAC1) were common to 2 out of 9 patients. Hence, for the majority of perturbed genes shared among LT survivors, no evidence was found about them being differentially expressed in a group comparison between LT and ST survivors. Among genes other than significant DEGs, only one was common to 7 out of 9 individuals: *NOSTRIN*, associated with nitric oxide pathways. No other genes were shared by 8 or all 9 LTS. Five out of 9 LTS patients shared *DTYMK* as perturbed gene in their individual transcriptome profile or PEEP. Six genes (*PDXDC1*, *ATF7IP2*, *LIN7C JTB*, *TTL*, *DVL2*), which regulate the ERG signal transduction pathways, were retained in 4 out of 9 LTS patients, were significantly involved in transcriptional mis-regulation in cancer (adjusted p-value=0.025). There were respectively 41 and 180 genes conserved, in 3 and 2 out of 9 LT survivors (Figure 5A; appendix pp 5: section 3.3).

**Figure 5:**
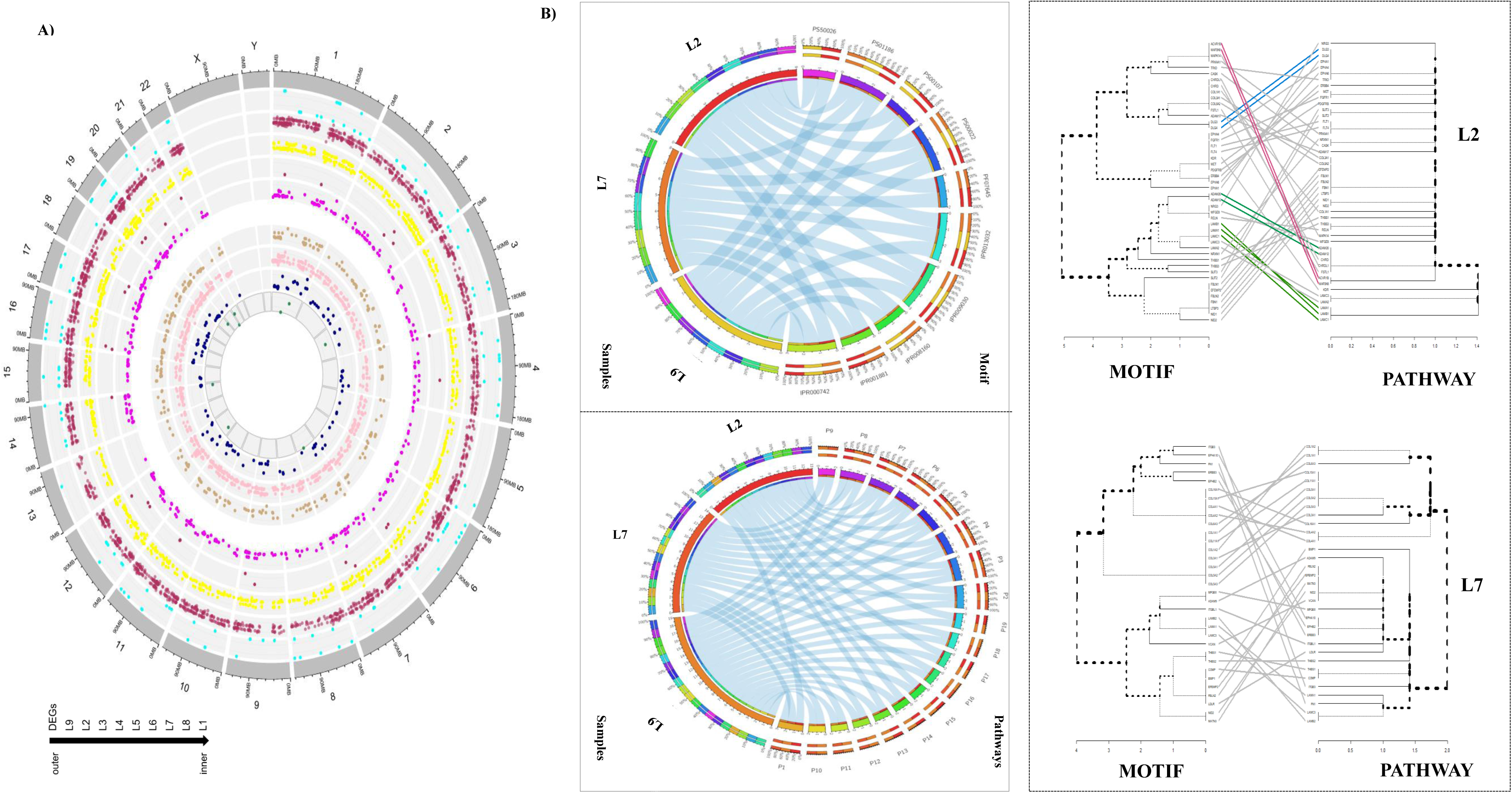
Genomic distributions of differentially expressed genes (DEGs) and PEEPs related to PDAC survivors using Circos plots and functional profiles of perturbation data: A) first outermost circle labeled with numbers represent chromosomes (same colors); the outermost track represents DEGs (up-regulated and down-regulated DEGs as scattered points); the nine innermost circles refer to the z-score for each LT survivor (LTS: ranging from LT1 to LT9) as scattered points. We have indicated perturbed genes only for chromosome 1 to 22 via track 2 to track 10 (outer to inner); B) Enriched KEGG pathways (P1 to P19 (out of 193)) and motifs common to at least 2 out of 9 LT individuals, shown via Circos Table Viewer (appendix pp 12-20). Each link refers to an LT survivor and a significantly enriched pathway (adjusted p-value < 0.05)/enriched motif based on the perturbed gene set found in that individual (data for LT2, LT7 and LT9 are shown). Uniquely enriched pathways across LT survivors are given in appendix pp 12-18; C) Visual comparison of two dendrograms developed from genes linked to enriched pathway and motif profiles. Similar sub-trees are connected with lines of the same color, while tree branches leading to distinct sub-trees are indicated with dashed lines.

### Individual-specific survival heterogeneity: Functional pathway and domain analysis in long-term PDAC survivors

We furthermore examined the extent to which the LT survivors reflected disruptions in KEGG and Reactome pathways, and identified pathways that were significantly enriched in at least one LT individual. In-depth analysis revealed 18 pathways in at least two LT survivors (Figure 5B; appendix pp 12-18). Thus, 175 pathways were uniquely perturbed in an LT PDAC survivor i.e. not shared among LT survivors (appendix pp 5: section 3.3).

Specific domain structures of genes play a significant role in gene regulation and expression. Hence, we also investigated the domain structures of perturbed genes in PEEPs of LTS to understand their potential regulatory mechanism in LT survival. A total of 47 enriched domains (adjusted p-value<0.05) were identified (appendix pp 19-20). The distribution of motifs that were commonly shared by 2 out of 9 LT survivors are shown in Figure 5B. For each LT survivor, we constructed two hierarchal trees based on the genes potentially involved in multiple domains and pathways, one for each for LT survivor. Resulting trees were statistically compared to identify the common branches. For instance, nine genes (indicated with pink, blue and green) were involved in common branches in LT2 (Figure 5C). More detail is given in appendix (pp 5: section 3.3).

### Exploitation of gene connectivity: systems views

Gene connectivity via reference networks can further highlight interesting gene clusters linked to LT survivors. In a first approach, we developed a disease module via DADA^5, 23^, which uses the human protein interactome network structure to prioritize disease genes, while at the same time removing possible biases induced by gene degree distributions (appendix pp 4). The disease module hypothesis proposes that disease regulatory genes should form one or a few large connected components in a human interactome. In this study, we restricted our seed genes (i.e., genes that play significant roles in PDAC according to the prior biological knowledge) to PDAC survival (*SMAD4*, *CDKA2*, and *KRAS*) and PDAC responsiveness based on a literature search and as identified from the DisGeNET database^24^ (appendix pp 21-22). Only the top 1% of DADA ranked genes were retained (Figure 6A I-IV; Figure 1:J), leading to 70 genes. Only one DADA top gene was also previously identified as DEG (*DKK4*), as shown in (Figure 6C). We also looked at the overlap between DADA-based 1% top-ranked genes and perturbed genes as highlighted by the PEEPs of individuals belonging to the long-term survival PDAC patient group (Figure 1:L). There were 23 genes in total. None of these common genes had previously been identified as DEGs. Out of 23, we identified 7 DADA top-ranked genes in common to clinical gene modules as identified before (Figure 6C; Figure 1:K; appendix pp 24-28). Only a single gene was shared by at least (actually exactly) three LT subjects, namely *GLI2*. Three genes (*RAC1*, *FOSL1*, and *EGF*) were shared by two out of 9 LT survivor PEEPs. Furthermore, three genes (*JAG2*, *TGFA*, *HDAC1*) were uniquely perturbed in a LT survivor (Figure 6B).

**Figure 6:**
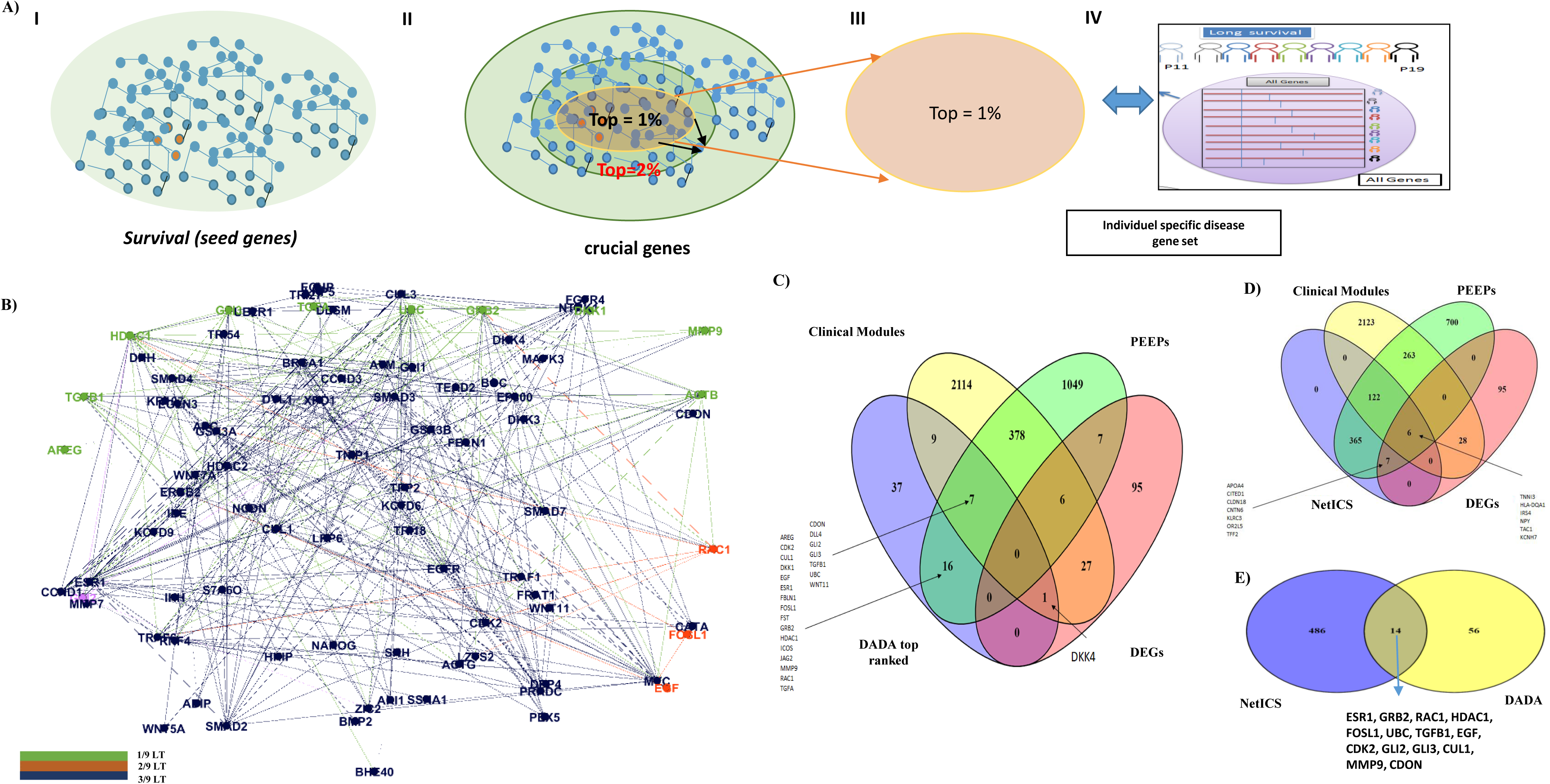
Exploitation of gene connectivity for LT PDAC survivor gene prioritization: A) DADA-oriented multi-step disease module identification: PDAC seed gene selection (I), restriction to top 1% of ranked genes (II-III) and intersection of retained gene list with individual perturbation gene expression profiles for LT survivors (IV); B) DADA-derived top-ranked genes found in at least one, two, or three LT survivors, indicated in green, orange and pink, respectively; C) Common genes to DADA and other gene prioritization approaches: DEGs, clinically relevant WGCNA gene modules, and PEEPs; D) Same as C) but with NetICS instead of DADA; E) Venn diagram showing the overlap between genes prioritized via NetICS and DADA. Common genes to top 1% NetICS individual gene lists and top 1% DADA genes are highlighted via arrows in C) and D).

In a second approach, we integrated individual-specific gene perturbation information (from PEEPs) with group-level DEG findings. For this, we used NetICS, which further allows unraveling inter- and intra-patient gene expression heterogeneity (appendix pp 4-5; Figure 1:E). Also in this approach a ranked list of genes was generated. The ranks are based on the gene scores, acquired through network diffusion algorithms (Figure 6D, appendix pp 23). Similar to the DADA approach, we focused on the top 1% of ranked genes for each LT survival patient, leading to 500 genes. Those 500 genes constituted a subset of PEEP genes. Only 13 genes out of 500 were also DEGs, including 6 genes that were additionally linked to clinical disease modules (Figure 6D). Among these 13 DEGs, *TNNI3* was NetICS top ranked, and was shared in its significance by 3 out of 9 LT survivors. It was also associated with the M7 module of clinical relevance (Figure 6D; Figure 1:G; appendix pp 24-28). Notably, *NOSTRIN*, a unique to NetICS gene (i.e., not highlighted by any other method shown in Figure 6D) was common to 7 out of 9 LT subjects. Furthermore, we found 14 genes common to DADA and NetICS gene prioritization methodologies (Figure 6E; Figure 1:J; appendix 40), involving the pathways such GPCR, Notch signaling pathway and many others. This common gene set did not include *TNNI3* nor *NOSTRIN*. The percentage of LTS PEEP genes not included in the top 1% DADA gene list is 27% (384/1440) and is similar to the percentage of LTS PEEP genes not included in the top 1% NetICS gene list (263/963).

## Discussion

Identifying molecular PDAC cancer drivers is critical for implementing precision medicine in clinical practice. Typically, the optimization and fine tuning of gene prioritization methods require large datasets^25^. Despite the small sample size of this study, we identified genes showing associations with multiple clinical traits^26^, and derived plausible links between long-term survival of patients and genes, pathways and protein domains by exploiting multiple approaches, including the combination of individual-level with group-level information in integrated analysis workflows. Throughout the entire study, we have relied on several statistical approaches to determine statistical significance with small samples.

PDAC accounts for over 90% of pancreatic cancer and is a lethal malignancy with very high mortality rates. The gene regulatory landscape of PDAC is defined by four mutational “mountains” (*KRAS*, *TP53*, *CDKN2A*, *SMAD4*), which are the main drivers of PDAC^27^. Cancer diseases are heterogeneous at different scales: group level, individual level, tumor type, cell level. This study reports on PDAC gene expression differences in patients who survived ≥36 months (long term LT) or <12 months (short term ST). Via advanced genomic profiling of PDAC survivors, we aimed to obtain more insights into LTS-relevant mechanisms that contribute to PDAC heterogeneity.

In this work, we identified known PDAC driver genes associated with survival, including *ROBO2, ZG16B*, and *PLXNA1*^28, 29^ (appendix pp 2). A thorough investigation of gene expression differences between LT and ST PDAC survivors highlighted gene involvement in immune responses (*CEACAM20*, *C6orf13*, *IRS4*, *CXCL17*), cell cycle (*SPDYE3*, *HLA-DQA2*, *CLDN*) and metabolic pathways (*GBA3*, *LIPN*), further highlighting the importance of these pathways in PDAC disease sruvival^30, 31^. All of these findings evidence that genes linked to immune responses could be useful in effective therapies for PDAC survival^32^. We also identified a downstream target of *KRAS* (*MUC16*) as DEG, supporting *KRAS* implications in survival^33^. Also, we observed modifications of *GKN1*, *KRT6*, and *ANKRD43* gene expressions in LTS, known to induce apoptosis and metastasis cancer^34, 35^. A previous study showed *REG4* as a serological marker for PDAC^36^. Very little information exists though about the role of *TSPAN8* in PDAC. However TSPAN8 promote cancer cell stemness via activation of Hedgehog signaling^37^. Furthermore, validation of DEGs via experimental work suggested a role of *REG4* and *TSPAN8* in PDAC survival mechanisms. Conserved domain database represents the curated information on conserved domain architectures of various proteins that shows implication in tumor initiation, tumor progression, angiogenesis and metastasis. The presence of multiple immunogenic domains (IGV, V-SET) in identified DEGs further supports recent activities towards cancer therapy^38^, and in-depth investigation of immunity cycles in relation to long-term survival in PDAC patients.

Systems biology approaches can provide immediate functional insights by revealing interactions between genes^39^. A motivation for WGCNA is that genes functioning together are regulated or co-expressed together^40^. Ballouz and cauthor^41^ suggested a minimal of 20 samples to predict meaningful functional connectivity. This forced us to pool STS and LTS together for WGCNA analysis on 19 patients and to link thus identified gene modules to clinical traits with non-parametric statistics whenever appropriate. Multiple studies have indicated an association of early survival in PDAC to tumor size^42^,. Additionally, multiple targets have been identified in the form of DEGs being associated with numerous traits such as tumor size, and the time between surgery and chemotherapy. In our study, identified several clinically relevant WGCNA gene modules (e.g., a gene module associated to time between surgery and chemotherapy with DEGs *LYZ, DKK4, CA14, NASE7, TSPAN8, GKN1, GKN2, SNORD116-18, DKK4*), which warrants further exploration on increased sample sizes in the future. Notably, *TSPAN8* serves as a prognostic marker in other cancer types as well^37^. Apart from time between surgery and chemotherapy, time to surgery may play an important role in PDAC as well (waiting for more than 30 days for surgery after diagnosis has been associated to an increase in tumor size)^43^. DEG *DKK4* (also top 1% DADA gene) is the least studied protein from the Dickkopf (*DKK*) family, which includes *DKK3*^44^ and *DKK1*^44^. The fact that *DKK4* did not appear in NetICS’s prioritization gene list, nor in PEEPs of LTS, seems to suggest that *DKK4* may be more promising in relation to controlling the survival of patients with PDAC rather than explaining individual heterogeneity among long-term PDAC survivors.

The identification of prognostic factors is complicated in the presence of individual-to-individual heterogeneity^45^. Detailed individual-specific omics profiling may be required to provide novel insights into LT survival in pancreatic cancer disease^46^. DEGs alone are unlikely to fully characterize individual (LT) survival, as observed for other complex traits^22^. Previous studies^14, 47, 48^ emphasized the existence of subgrouping of PDAC patients in general, based on expression profiling of samples. Our study showed that any LTS patient only exhibits a small fraction of group-wise DEGs in their PEEP profiles and shows a deep level of gene expression heterogeneity. Notably, several genes were uniquely perturbed in an LT survivor, which strengthens our belief that LTS patients exhibit more abundant levels of heterogeneity. Careful inspection of PEEPs across LT survivors highlighted biological signatures: focal adhesion^49^, and ECM receptors^50^. Interestingly multiple PDAC responsive pathways^51^ were enriched across several LT survivors and led to further subgrouping of LT survivors. Understanding these pathways may provide novel insight into the LT survival mechanism in PDAC. PEEP analysis identified *FCGR3A*, a potential biomarker in PDAC^52^. Two genes, *NOSTRIN* and *ADGRG6*, were shared by 66% of LTS, and have been reported before to be associated with PDAC survival.^40, 53^

Drugs bind to their target proteins and perturb the transcriptome of a cancer cell^54^. In our study, analytic functional analysis of individual PEEPs helped to decode homogeneity patterns within LTS. Heterogeneity at the gene level may go hand in hand with homogeneity at the pathway level as different gene perturbations may lead to disruptions in the same molecular pathway. The use of network-centric approaches resulted in various oncogenes such as *CUL1*, a central component of *SCF*^55^, *EGF*, *FOSL1*^56^, *MMP9*^57^, and *TGFB1*^31^, already known as anticancer targets. Different transcription factors (*GLI2* and *GL3*) were identified, linked to the *KRAS* mechanism of pancreatic tumorigenesis^58^. Identified Immunogenic gene (*CDON*) and regulatory gene (*HDAC1*) targets could play significant roles in the future immunotherapeutic strategies in long-term PDAC survivors^46^. *CD8* revealed in our study is in line with recent studies in which *CD8* expression profiling was linked to an immunologic subtype of PDAC with favorable survival^59^. These results indicate the advantages of adopting integrative analysis pipeline that combining knowledge about network-driven disease modules with individual-specific gene perturbation profiling even for small sample sizes,. Unlike DEG-oriented therapeutic target selection for cancers, commonly used to date, we promote the exploitation of analytic frameworks in which multiple network-centric approaches are used for the identification of patient-specific therapeutic targets. This will boost cancer prognosis and treatment in the context of personalized medicine.

## Conclusion

For the first time in PDAC patients, we demonstrated and applied an integrative analytic workflow that combines clinical and omics data, as well as individual- and group-level transcriptome profiling. We showed the utility of network-based approaches, disease modules and multi-scale functional analyses (gene, protein domain, pathway), leading to the identification of known oncogenes and previously unreported marks contributing to heterogeneity in long-term PDAC survival.

## Contributions

A.B under supervision of K.V.S performed the detailed BIOINFORMATICS data analyses. D.V.D. and M.C. designed the protocol to obtain patient materials and supervised the extraction of clinical information from CHU Liège databases. A.B. and K.V.S wrote the manuscript with input from all co-authors. C.J performed the nucleic acid extractions. C.J, I.S and C.P participated in the scientific discussions related to the conception of this manuscript.

## Declaration of Interests

The authors declare no competing interests

## Acknowledgments

We thank the Biobanque of Liège University Hospital and the GIGA Genomics Platform for sequencing. A.B, I.S and K.V.S acknowledge funding by Télévie 2015 “PDAC-xome: Exome sequencing in PDAC” (convention n° 7.4629.15), Télévie 2016 “Drivers and markers in pancreatic cancer” (convention n° 7.4502.16), and FRS-FNRS – CDR 2017 “SysMedPC” (convention n° J.0061.17).

## Accession Numbers

Data deposited in GEO with accession number GSE150043.

## Appendix Method

## Data and Methods

### 1. Data and data preparation

#### 1.1 Patient selection, ethical statement, and criteria to maximize the definition of STS and LTS

All aspects of the study comply with the Declaration of Helsinki. PDAC patients from Liege University Hospital were recruited on the basis of an opt-out methodology, from 2007 to 2014, giving to N=96 pancreas tissue. Tissues were obtained from the University of liege Biobank, Belgium. The study was approved by the local institutional ethical board (“Comité d’éthique hospital-faculties universities de Liège (707)) under the file number B707201627153. Among them, 36 had OS < 12 months or > 36 months, as selected survival criteria. We performed RNA extraction from those 36 samples and processed for RNA quality check.

#### 1.2 RNA extraction, library preparation, sequencing

Tumor areas were determined by a certified pathologist and were manually macro-dissected from the FFPE tissues. RNA was extracted using an All Prep DNA/RNA/miRNA Universal kit (Qiagen, Belgium) according to the manufacturer’s protocol. The next manipulations described in the paragraphs were performed by the GIGA-Genomics facility. The RNA quality (N=36) was assessed using a BioAnalyzer (Agilent, Belgium), and the proportion of RNA with a length higher than 200 bases (DV200) was measured. Only 19 out of 36 met a suitable RNA quality, allowed for sequencing. TruSeq® RNA Access Library Prep Kit (Cat. No. RS-301-2001 and RS-301-2002) (Illumina, The Netherlands) was used to prepare libraries, and next-generation sequencing was performed on a NextSeq500 apparatus (Illumina, The Netherlands), in paired-end 2 x 75bp high output mode.

We performed a series of transcriptome computational analyses to better understand patient heterogeneity between LT and ST survivors. After quality control and adaptor trimming with Trimmomatic^1^, sequence data were mapped to the Genome Reference Consortium GRCh38 assembly using STAR v2.5.2^2^. Read counts for known genes were generated using the function HTSeq-count v0.6.1p^3^ and data were normalized in DESeq2 v1.20.0^4^ as shown in Figure 1.

#### 1.3 Clinical features of Patients

Various clinical and pathological parameters of patients (N=19) were included in the analysis. In particular, we collected the following pathological clinical data: age, sex, tumor size, number of lymph nodes evaluated, tumor grade, sugey magins invaded by tumor cells, time between surgery and chemotherapy (in days), time between surgery and relapse (in months), disease-free survival (DFS), vascular resection, time in hospital after surgery (in days), re-hospitalization 6 months after surgery, vascular contact, artery contact, and chemotherapy as shown in Figure 2A.

#### 1.4 Group based DEGs analysis: Differential Gene analysis and functional follow-up

We used DEseq2^4^ for the identification of differentially expressed genes (DEG), with the thresholds log2 fold change ≥2 and ≤−2, to indicate up-regulation and down-regulation, respectively (Figure 1:B). Significance was assessed at an unadjusted p-value <0.05 in LT vs. ST group comparison^5^. We used the ClusterProfiler v3.8.1^6^ package to predict various GO processes enriched in differentially expressed genes (DEGs). To identify the protein domain in DEGs, we used batch CD-Search^7^. Identified DEG were analyzed for detection of survival genes, with a log-rank test in a Kaplan–Meier survival model^8^ (Figure 1:A-B). For each gene, patients were classified into two groups, the high-expression group (H) and the low-expression group (L), using the expression median of the gene as a cutoff using the survminer^9^ (v. 0.4.6) R package.

### 2. Methods

The entire workflow is described in Figure 1. Specific details regarding group-level and individual-specific analyses are given in Sections 2.1 and 2.2, respectively.

#### 2.1 Group-level survival heterogeneity: WGCNA for gene module prediction and assessment of clinical relevance

The minimum sample size to run weighted gene co-expression network analysis (WGCNA) is at least 15. Therefore, WGCNA v1.63^10^ was used on pooled ST and LT PDAC survival patients to generate a transcriptional network from the normalized expression data. The weighted coefficient β was selected based on scale-free topology criteria. The adjacency coefficient α was computed using the power to measure the correlation strength between two genes. The adjacency matrix was created based on α, which was subsequently transformed into a topological overlap matrix (TOM). The distance measure dissTOM = 1−TOM, served as input to perform average linkage hierarchical clustering (with DynamicTreeCut^11^), giving rise to gene co-expression modules. Gene modules were shown as branches of the resulting pruned tree. It was followed by the calculation of module eigengenes (MEs), which are defined as the 1st linear principal component of each co-expression module. The hierarchical clustering of MEs was performed to study associations between modules. Approximate non-parametric association tests were used to investigate the association between MEs and PDAC clinical traits. In effect, we used two methods to identify modules related to clinical progression traits. First, within-module gene significance was identified for every module and all available clinical traits. Average gene significance for a module was defined as “module significance”, following recommendations of ^12^. Second, rank-based correlation (r) was performed among each ME with the multiple clinic pathological characteristics available in this study (adjusted p-value for 0.05 MEs). We used parametric (Pearson correlation coefficient) and non-parametric (Spearman rank) tests for each continuous and categorical data, respectively. In order to assess the functional relevance of clinically associated modules, we used ClueGO^13^, a Cytoscape plug-in in order to visualize the non-redundant biological terms for genes in a functionally comparative network from multiple clusters. Non-redundancy was assessed via two-sided hypergeometric testing for enrichment/depletion (Bonferroni adjusted p-value < 0.05). Cytoscape 5.0^14^ was used for visualizing gene interaction networks (Figure 1:C).

#### 2.2 Individual-specific survival heterogeneity: Quantification of heterogeneity between individual transcriptome profiles, with functional and clinical relevance

We used principles of the PEPPER^15^ method to construct personalized gene expression perturbation profiles for each of N=19 PDAC subjects. PEPPER requires a target class of individuals and a reference class (Figure 1:D). In this study, we took LT PDAC survivors as target group and considered ST survivors as reference (i.e., the most abundant group in real-life). The approach captures the extent to which gene i is perturbed in subject j via a Z-score. This Z-score indicates how many standard deviations the individual’s gene expression is away from the mean value of the reference group. As a threshold, we used |z| =2. Positive z-scores > 2 would indicate up-regulation, negative z-scores < −2 would indicate down-regulation. Given the small sample sizes to work with in this study, we reshuffled the ST/LT group labels^16^ 500 times, and repeated the experiment. Note that under the null hypothesis, none of the individual LT survivor profiles would be markedly different from average ST survivor profiles and thus LT/ST survivor status would be exchangeable on the basis of individual transcriptome profiles. We used shinyCircos^17^ R package to develop circos plot for identified PEEPs. Functional follow-up analyses included checking for enrichment of KEGG pathways, and verifying motif enrichment via ToppGene Suite^18^ (multiple testing adjusted p-value < 0.05). Also, patient-specific one-way hierarchical clustering and dendrograms were developed on the basis of the frequency of perturbed genes in identified domains and pathways. Both dendrograms were subsequently compared using the R version 1.12.0 of the dendextend^19^ R package”. For deeper insights, two-way clustering via the superbiclust package in R (RcmdrPlugin.BiclustGUI^20^) version 1.1) was used, enabling the application of the Bimax^21^ algorithm to jointly cluster LT survivors and either one of three levels of biological information, namely gene, pathway and motif levels. For each analysis, a higher level (super) biclustering was obtained by constructing a hierarchical tree depicting Jaccard similarity between Bimax clusters.

In the aforementioned PEEPs analyses (PEEP: an individual perturbation expression profile against a reference), no notion of gene-connectivity was used. However, gene connectivity via reference networks can further highlight interesting gene clusters linked to LT survivors. Here, we considered physical interaction data as available from ConsensusPathDB^22^, and obtained 373,101 links between N=19,117 genes. Starting with genes in pathways that already have been implied in PDAC via ^23^, and supplementing these genes with searches in the DisGeNet database^24^ (search term = “Pancreatic Diseases”), resulted in 53 seed genes (Figure 1:I; appendix pp 19-20). We then used DADA’s module detection algorithm^13^ to augment the initial list of 53 seed genes and to identify PDAC disease modules. The top 1 percent highest ranked genes were considered to form a disease module. Significantly perturbed genes (in LT survivor PEEPs) were mapped on the identified disease module. This allowed putting LT survival individual specific genes in the context of gene connectivity and gene neighborhoods. All DADA top 1 percent genes were checked for their retrieval in previous analyses (Figure 1:J-L). As an alternative approach to exploit gene interaction network structure, we adapted NetICS^25^, an approach initially intended to prioritize cancer genes on a directed functional interaction network. It uses an individual-specific list of genes via bidirectional network diffusion of two layers of information (Figure 1:E). As first layer we took the individual-specific significant genes as highlighted in the LT PDAC survival PEEPs analyses before (instead of mutant genes per sample in the original NetICS implementation). As second layer we took groups-specific DEGs (Section 2.1). Individual-specific gene ranks (for LT survivors) were aggregated via NetICS methodology into an overall ranked list of genes, with restart probability of 0.4. The top 1% percent ranked genes were retained. Similar to follow-up of DADA top-ranked genes, we checked for the frequency of NetICS derived top-ranked genes that were also retrieved in former analyses (Figure 1: F-H).

### 3. Results

#### 3.1 Potential candidate genes

Gene *XKR5,* showed a significant increase in survival in long-term (LT) patients with lower expression compared to short-term survivors (appendix pp 27). We observed a similar pattern for the genes *GATD3B*, *CYP27C1,* and *miR-765* (appendix pp 28-30). These results highlight the potential of the identified genes in further understanding molecular underpinnings of PDAC survival.

#### 3.2 Functional analysis of clinically relevant modules

Five modules were identified as clinical relevant module via WGCNA analysis (appendix 3). In module M34, we found three significant Reactome pathways distributed into three groups: the effects of PIP2 hydrolysis (4 genes), the deactivation of the beta-catenin transactivating complex (3 genes) and the VEGFA-VEGFR2 pathway (4 genes) (data not shown). In M30, we found two significant pathways: apoptotic cleavage of cell adhesion proteins (4 genes) and o-linked glycosylation (11 genes) (data not shown). Individuals (LT1, LT3, LT4, LT5, LT6) did not show significant enrichment in any KEGG/Reactome pathway.

#### 3.3 Biclustering of functional profiles

To analyses the effect of perturbed genes in PEEPs (gene is significantly perturbed or not) in LT survivor’s heterogeneity, two-way clustering (biclustering) highlighted 64 gene clusters (appendix pp 33-34). The largest cluster (cluster 15) consisted of 363 genes. Deeper hierarchical clustering of these identified clusters (appendix pp 4) grouped cluster 7, 36, 37, 42, 47,48, 50, 53, 55 into single super cluster (appendix pp 34) with overrepresentation of cancer specific pathways such as mTOR pathways and NOD-like signaling pathways as highlighted via orange box in appendix (pp 36).

Two-way hierarchical clustering, based on the presence/absence of enriched pathways across LT survivors (LT2, LT7, LT8, LT9) revealed three clusters (appendix pp 38). First two clusters (C1 and C2) showed enriched pathways in two LTS only. C1 consisted of 14 pathways was collectively enriched in L7 and L9, and highlighted a strong association with cancer-related pathways. C2 showed enrichment of 13 pathways between L9 and L2 such as Proteoglycans in cancer and EPH-Ephrin signaling. Smallest cluster C3 consisted of 8 pathways across three LTS survivors i.e LT2, LT7, LT9. Deeper hierarchical clustering groups C2 and C3 into single supercluster based on similar pathways profiles. Likewise, two-way hierarchical clustering (biclustering) based on motif enrichment profiles (present or absent) across all LT survivors resulted in four clusters (appendix pp 39). The first cluster (C1), represented by LT7 and LT9, was enriched with six domains. The second cluster (C2), active in LT2 and LT7, was enriched with 7 domains. The third clusters (C3) involved enrichment of 7 domains shared two among LT survivors (appendix pp 17-18). The fourth cluster (C4) was largely shared by three LT survivors (LT2, LT7, and LT9). This cluster involved 5 domains: IPR013032, PS01186, IPR000742, PS00022, and IPR009030. Deeper hierarchical clustering groups C1 and C4 into single supercluster based on similar protein domains profiles. Deeper hierarchical clustering groups C1 and C4 into single supercluster based on similar protein domains profiles. More in-depth analysis revealed a common gene set between cluster 24 obtained from gene-level clustering and cluster 1 (C1) derived from pathway-level biclustering (appendix pp 36, 39). Similarly, cluster 25 derived from gene level analysis showed overlap with cluster 2 (C2) derived from pathway-level biclustering.

---

**Table S1.** Complete list of all sample (patients) and their clinical features as considered in the present study.

| Patient ID | Group* | Age at the surgery (years) | Sex<sup>a</sup> | Tumor size (cm)<sup>b</sup> (measured after surgery) | T | N | Number of nodes analyzed |
| --- | --- | --- | --- | --- | --- | --- | --- |
| S1 | ST | 74 | F | 3 | 4 | 1 | 25 |
| S2 | ST | 71 | M | 3.5 | 3 | 0 | 12 |
| S3 | ST | 72 | F | 3.2 | 3 | 0 | 7 |
| S4 | ST | 73 | M | 4 | 3 | 1 | 7 |
| S5 | ST | 54 | F | 3.2 | 4 | 1 | 12 |
| S6 | ST | 80 | M | 4 | 3 | 1 | 8 |
| S7 | ST | 80 | M | 4 | 3 | 0 | 16 |
| S8 | ST | 63 | F | 3 | 4 | 1 | 14 |
| S9 | ST | 66 | M | 5 | 3 | 1B | 16 |
| S10 | ST | 66 | M | 5 | 4 | 1 | 4 |
| L1 | LT | 78 | F | 2.5 | 3 | 1 | 13 |
| L2 | LT | 58 | M | 3.2 | 3 | 0 | 33 |
| L3 | LT | 71 | F | 2 | 3 | 0 | 10 |
| L4 | LT | 83 | M | 4 | 3 | 0 | 14 |
| L5 | LT | 67 | M | 2.2 | 3 | 1 | 25 |
| L6 | LT | 62 | M | 5 | 3 | 1 | 7 |
| L7 | LT | 58 | M | 2 | 1 | 1 | 8 |
| L8 | LT | 77 | M | 4.5 | 4 | 1b | 4 |
| L9 | LT | 55 | M | 0.6 | 1 | 0 | 0 |

Complete list of all sample (patients) and their clinical features analyzed in the present study (Table S1 continue)
| Patient ID | ratio node<br>(invaded/analyzed) | ratio node %<br>(invaded/analysed) | Sugey Magins<br>invaded by<br>tumor cells** | Time between<br>surgery and<br>chemotherapy<br>(in days) | Time between<br>surgery and relapse<br>(months) (DFS)<sup>c</sup> | Time between<br>surgery and<br>death (months)<br>OS<sup>d</sup> | Time between<br>relapse and<br>death<br>(months) | Death | Vascular<br>resection |
| --- | --- | --- | --- | --- | --- | --- | --- | --- | --- |
| S1 | 2/25 | 0.08 | 1 | 82 | NA | 3.09 | NA | 1 | 1 |
| S2 | 0/12 | 0.00 | 0 | NA | 5.00 | 6.31 | 1.31 | 1 | 1 |
| S3 | 0/7 | 0.00 | 0 | 71 | NA | 7.23 | NA | 1 | 0 |
| S4 | 5/7 | 0.71 | 0 | 54 | 5.42 | 8.32 | 2.89 | 1 | 0 |
| S5 | 7/12 | 0.58 | 0 | 63 | 2.30 | 8.42 | 6.11 | 1 | 0 |
| S6 | 2/8 | 0.25 | 1 | 74 | 4.50 | 8.61 | 4.11 | 1 | 0 |
| S7 | 0/16 | 0.00 | 0 | NA | 6.87 | 9.66 | 2.79 | 1 | 0 |
| S8 | 4/14 | 0.28 | 1 | 43 | 4.87 | 9.66 | 4.80 | 1 | 1 |
| S9 | 10/16 | 0.62 | 0 | 53 | 7.10 | 9.76 | 2.66 | 1 | 0 |
| S10 | 0/4 | 0.00 | 0 | 41 | 9.37 | 10.22 | 0.85 | 1 | 0 |
| L1 | 4/13 | 0.31 | 0 | 55 | 25.25 | NA | NA | 1 | 0 |
| L2 | 0/33 | 0.00 | 2 | NA | 17.16 | 40.70 | 23.54 | 1 | 1 |
| L3 | 0/10 | 0.00 | 1 | NA | NA | NA | NA | 1 | 0 |
| L4 | 0/14 | 0.00 | 0 | NA | NA | 48.72 | NA | 1 | 0 |
| L5 | 2/25 | 0.08 | 0 | 16 | NA | NA | NA | 0 | 0 |
| L6 | 1/7 | 0.14 | 0 | 71 | 37.71 | 60.52 | 22.81 | 1 | 0 |
| L7 | 3/8 | 0.37 | 0 | NA | NA | NA | NA | 0 | 0 |
| L8 | 3/4 | 0.75 | 0 | 66 | NA | NA | NA | 0 | 0 |
| L9 | 0 | NA | 0 | NA | NA | NA | NA | 0 | 0 |

Complete list of all sample (patients) and their clinical features analyzed in the present study (Table S1 continue)
| Patient ID | Time in hospital after surgery (in days) | Re-hospitalisation 6 months after surgery** | Vascular contact (if tumor contact a vein (often portal vein)** | artery contact (if tumor contact an artery) ** | Tumor size by imagery (mm)<sup>e</sup> | Time between imagery and surgery (days) | Chemotherapy (CTH)** | CTH > 3 months | Number of CTH cures | Dose CTH received in % of the maximal theoretical value |
| --- | --- | --- | --- | --- | --- | --- | --- | --- | --- | --- |
| S1 | 42 | 0 | 1 | 0 | 42 | 7 | 1 | 0 | 1 | 100 |
| S2 | 14 | 0 | 0 | 0 | 76 | 121 | 0 | NA | NA | NA |
| S3 | 16 | 0 | NA | NA | NA | NA | 1 | 0 | NA | NA |
| S4 | 22 | 3 | 1 | 0 | 22 | 6 | 1 | 0 | 1 | 100 |
| S5 | 16 | 0 | 1 | 0 | 29 | 6 | 1 | 0 | 4 | 100 |
| S6 | 24 | 0 | 1 | 1 | 34 | 5 | 1 | 0 | 6 | 80 |
| S7 | 21 | 1 | 1 | 1 | 27 | 3 | 0 | NA | NA | NA |
| S8 | 18 | 0 | 1 | 0 | NA | 31 | 1 | 0 | 7 | 100 |
| S9 | 24 | 0 | 1 | 1 | 19 | 21 | 1 | 0 | 3 | 100 |
| S10 | 17 | 0 | 1 | 0 | NA | 92 | 1 | 0 | 2 | NA |
| L1 | 22 | 0 | 1 | 0 | 15 | 9 | 1 | 0 | 4 | 100 |
| L2 | 14 | 1 | NA | NA | NA | NA | 1 | 1 | NA | NA |
| L3 | 11 | 0 | 1 | 0 | 28 | 22 | 0 | NA | NA | NA |
| L4 | 22 | 0 | 1 | 0 | NA | 27 | 0 | NA | NA | NA |
| L5 | 17 | 0 | 1 | 0 | 28 | 33 | 1 | 0 | NA | NA |
| L6 | 17 | 0 | 0 | 0 | NA | 63 | 1 | 0 | 4 | 75 |
| L7 | 13 | 0 | 1 | 1 | 21 | 28 | NA | NA | NA | NA |
| L8 | 28 | 0 | 1 | 0 | 41 | 28 | 1 | 0 | 18 | 100 |
| L9 | 21 | 0 | 1 | 0 | 21 | 28 | 0 | NA | NA | NA |

\* ST=short-term PDAC survivor (ST; death $\geq 3$ months et $< 12$ months) ; LT= long-term PDAC survivor (LT; alive or death $\geq 36$ months) ; <sup>a</sup>F= Female ; M=Male ; <sup>b</sup>cm = centimeters ; <sup>c</sup>DFS=Disease free survival ; <sup>d</sup>OS= Overall survival ; <sup>e</sup>mm= millimeters ; NA =Not available; \*\* (1=yes;0=no)

**Table S2.**
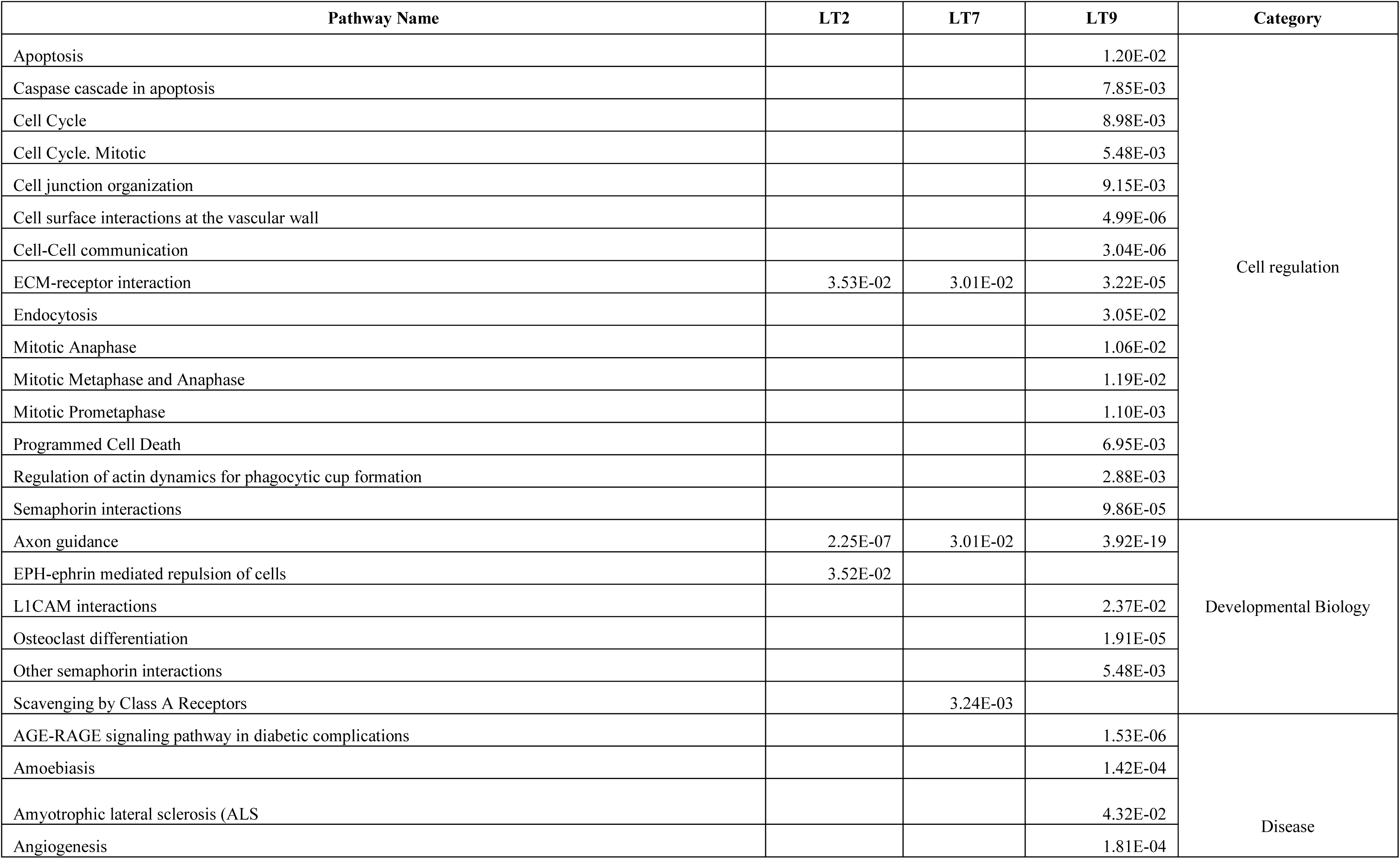
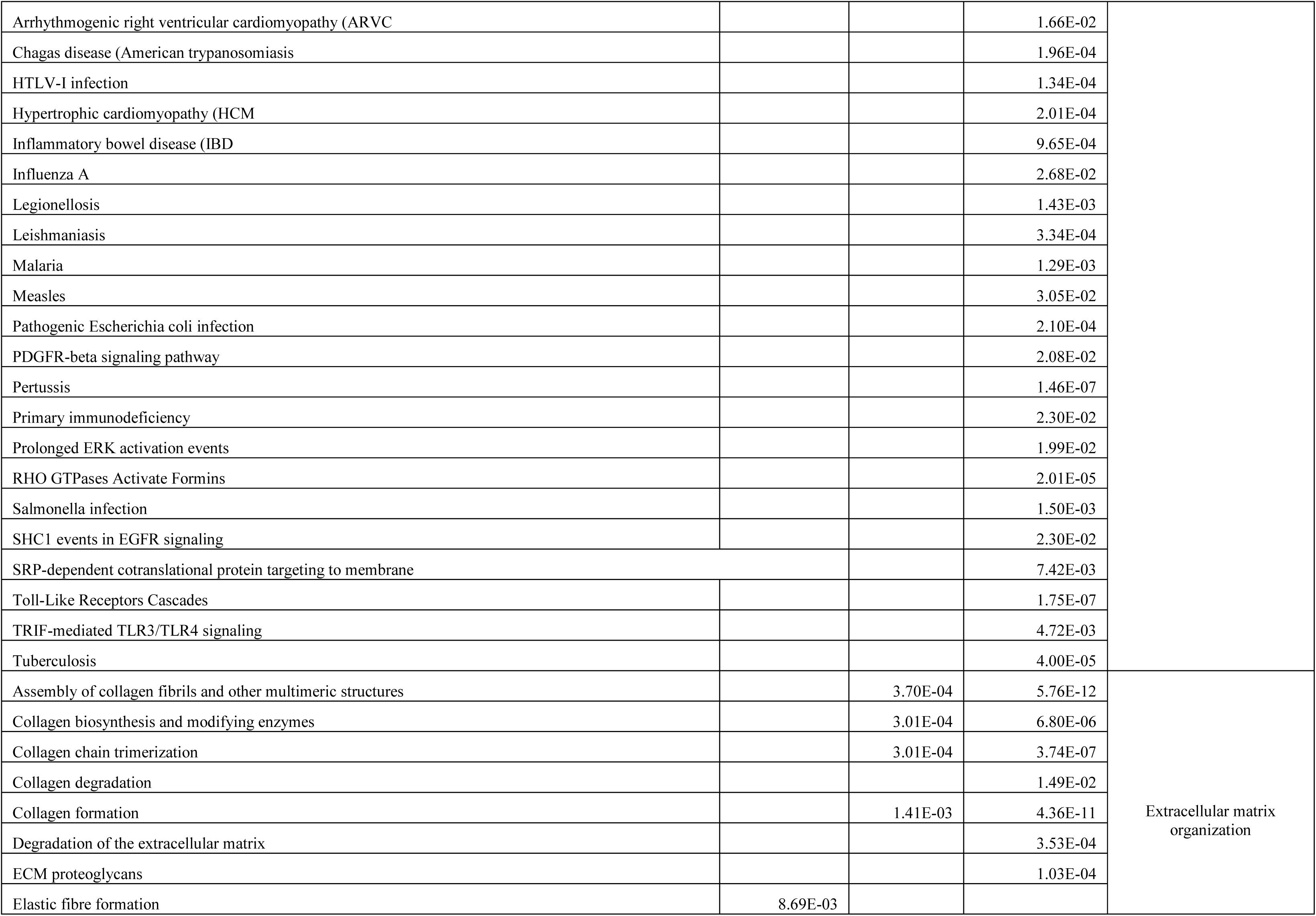
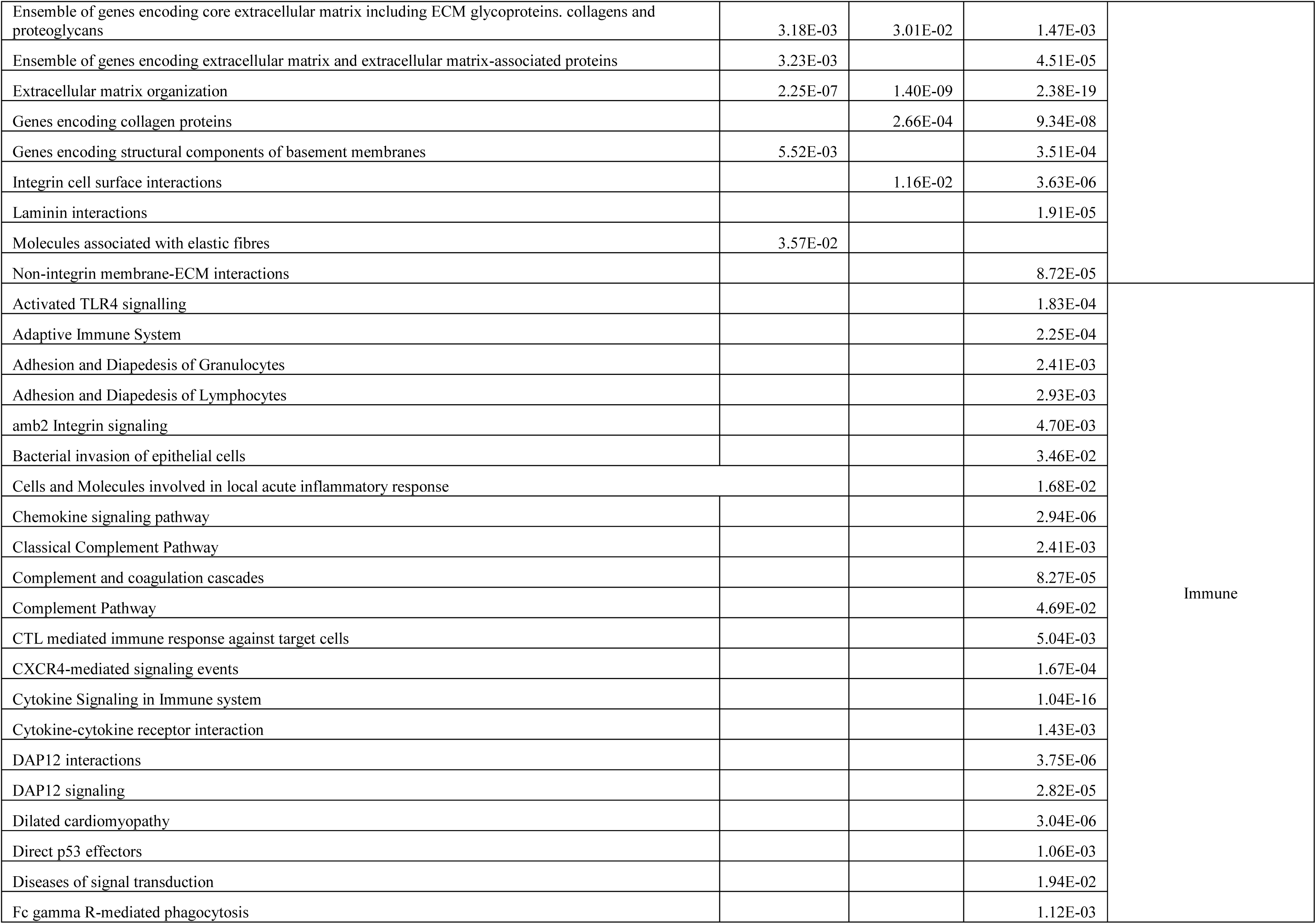
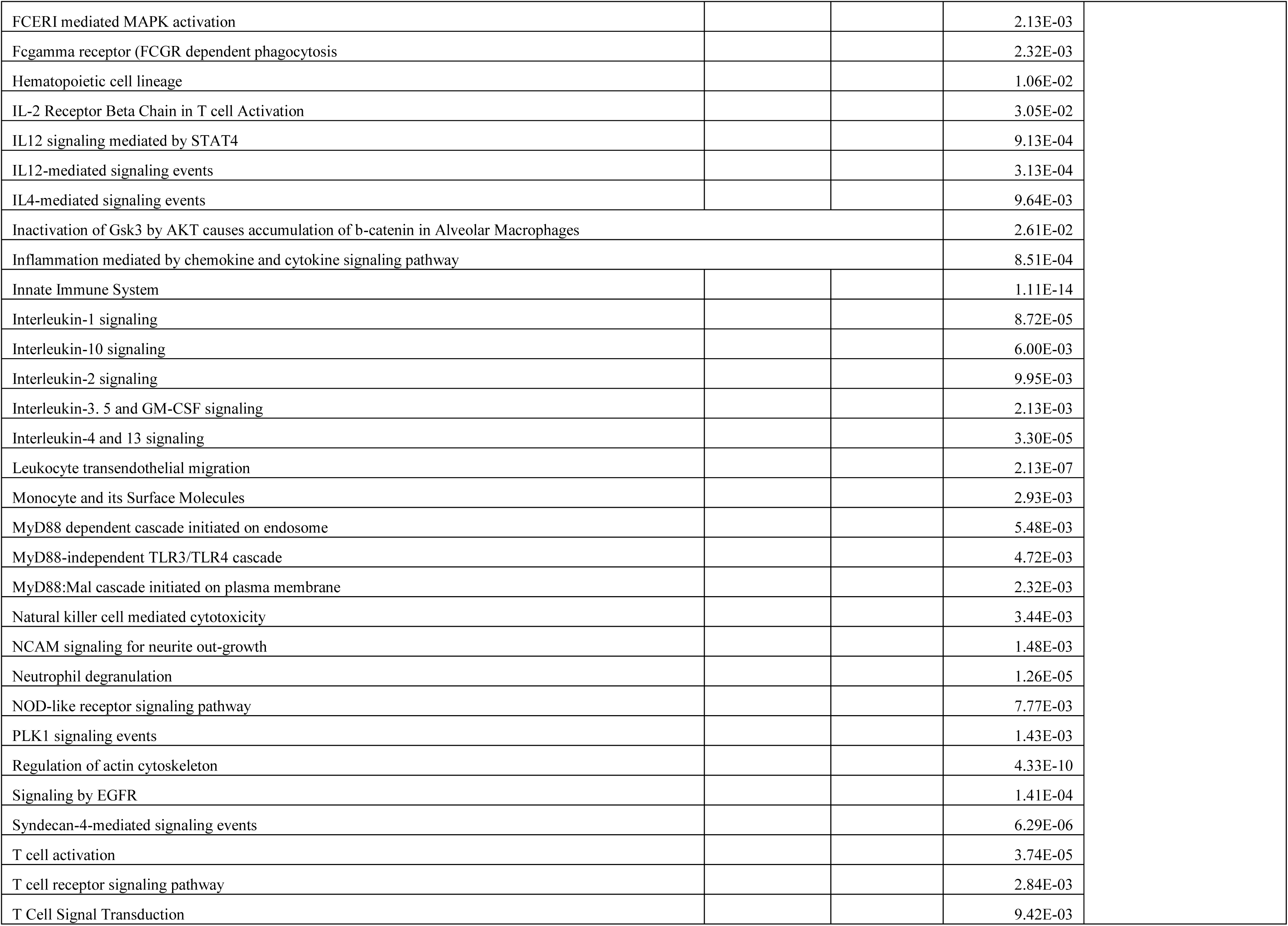
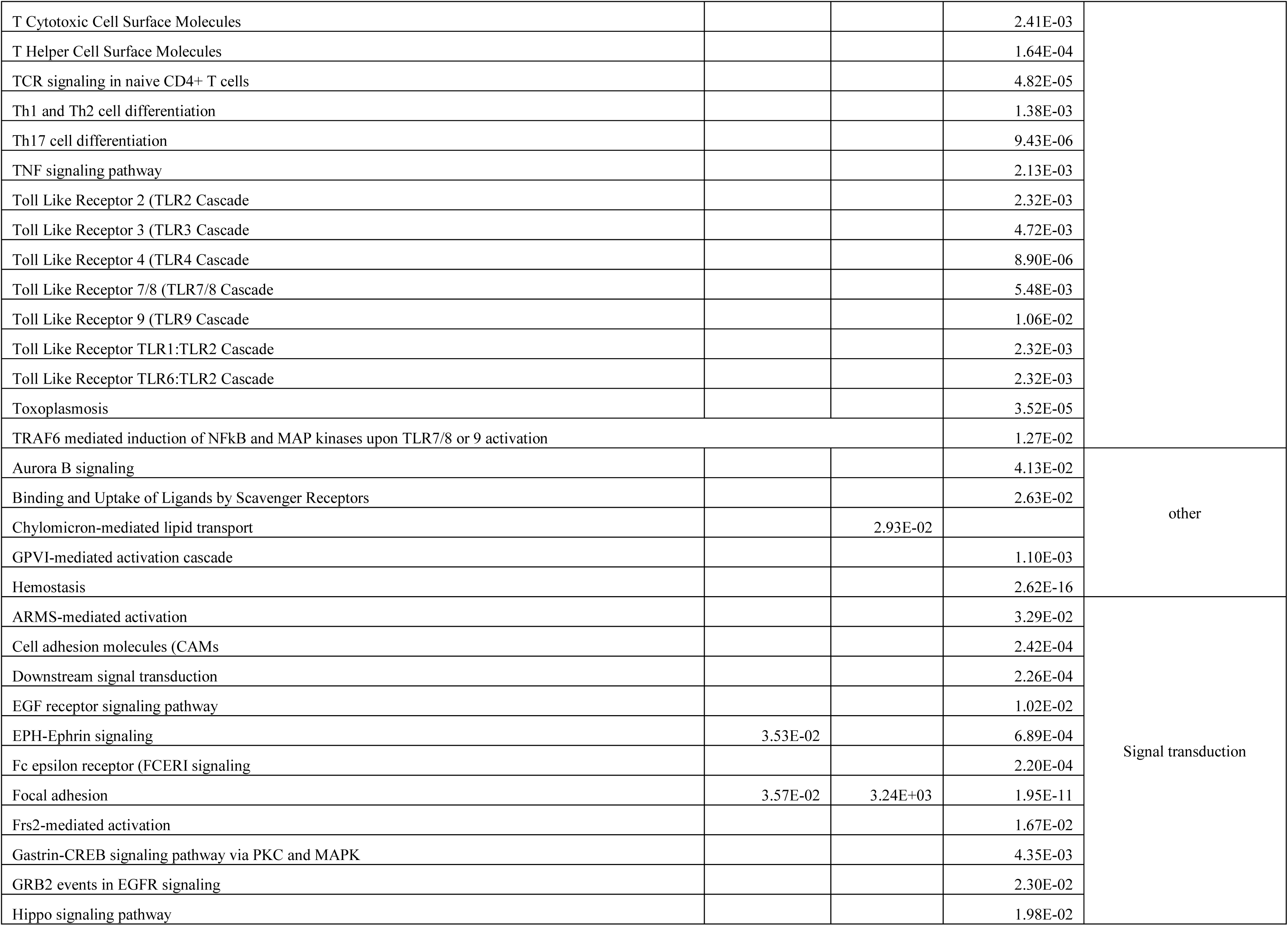
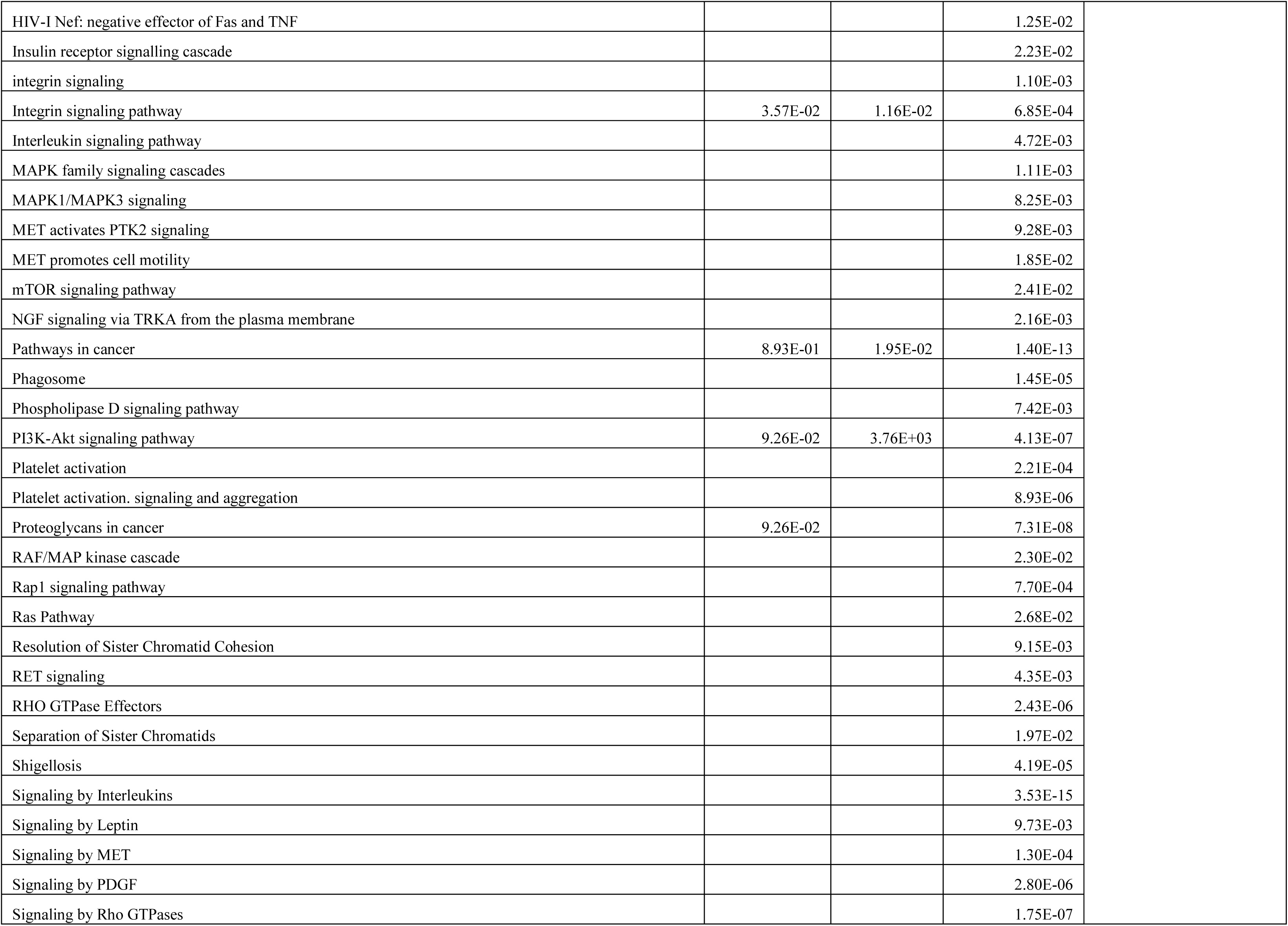
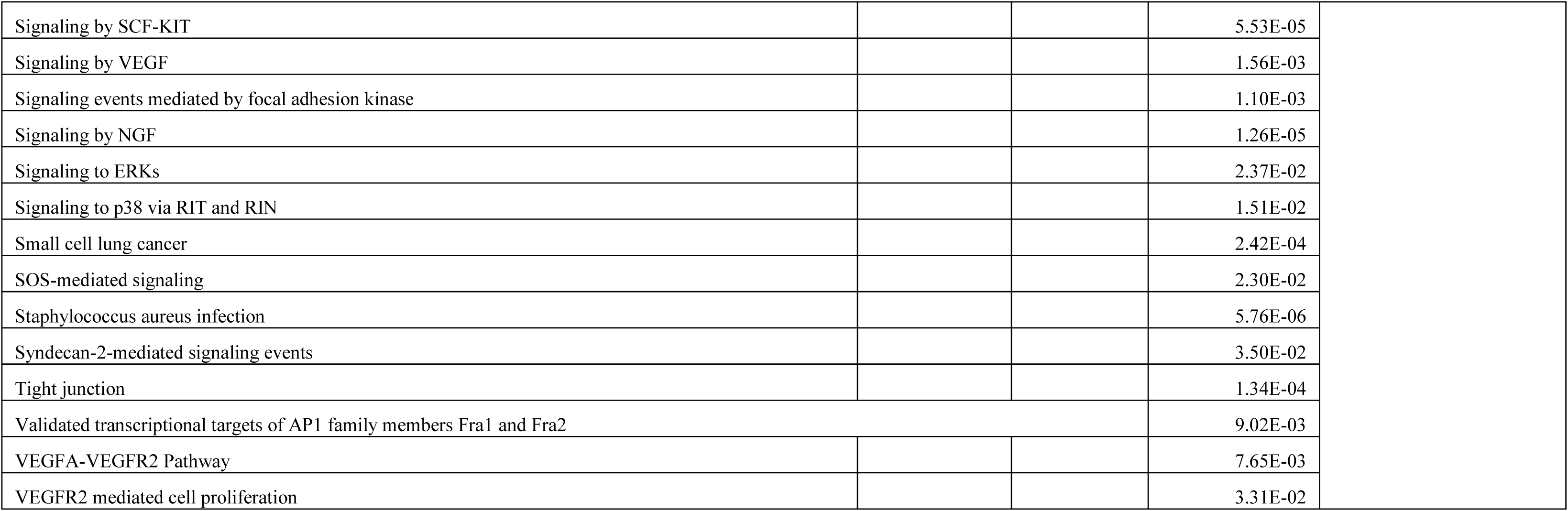
**Pathways enriched in LT survivors based on the Individual specific PEEPs annotated using ToppGene Suite**^17^ **(section 2.2).** For all identified significant pathways, corresponding Bonferroni corrected p-values are shown. Identified pathways are grouped into various categories based on functional annotation. highlighted based on their involvement either in three LT survivors; two LT survivors or one LT survivor.

**Table S3.** **Protein domains enriched in LT survivors based on the Individual specific PEEPs annotated using ToppGene Suite**^17^ **(section 2.2).** For all identified significant protein domains, corresponding Bonferroni corrected p-values are shown. Identified protein domains are ordered based on similar annotation profile as given below. (Only 3 LT survivors showed enriched IPR domains).

| Domain ID | LT2 | LT7 | LT9 | Description |
| --- | --- | --- | --- | --- |
| PS00010 | 0.000368 |  |  | ASX_HYDROXYL |
| PF12662 | 0.000247 |  |  | cEGF |
| IPR026823 | 0.000247 |  |  | cEGF |
| SM00038 |  | 2.5E-05 |  | COLFI |
| PF01410 |  | 2.5E-05 |  | COLFI |
| PF01391 |  | 0.001874 | 4.19E-06 | Collagen |
| IPR008160 |  | 0.001874 | 4.19E-06 | Collagen |
| IPR011029 |  |  | 0.00017 | DEATH-like_dom |
| SM00181 | 2.1E-06 | 7.05E-05 | 1.5E-06 | EGF |
| PF00008 |  |  | 9.42E-06 | EGF |
| PS00022 | 8.89E-05 | 5.77E-05 | 4.19E-06 | EGF |
| PS01186 | 9.45E-06 | 4.21E-06 | 7.02E-06 | EGF |
| PS50026 | 0.000011 |  | 0.000209 | EGF |
| PF07645 | 5.51E-05 | 0.000249 |  | EGF |
| SM00179 | 2.47E-05 | 0.001874 |  | EGF |
| PS01187 | 5.01E-05 |  |  | EGF |
| IPR018097 | 4.01E-05 |  |  | EGF_Ca-bd_CS |
| IPR001881 | 2.9E-05 | 0.002076 |  | EGF-like_Ca-bd_dom |
| IPR013032 | 2.16E-06 | 1.02E-05 | 4.79E-08 | EGF-like_CS |
| IPR000742 | 9.45E-06 | 4.12E-05 | 7.07E-04 | EGF-like_dom |
| IPR000152 | 0.000121 |  |  | EGF-type_Asp/Asn_hydroxyl_site |
| PD002078 |  | 2.5E-05 |  | Fib_collagen_C |
| IPR000885 |  | 2.5E-05 |  | Fib_collagen_C |
| SM00060 |  |  | 0.000799 | FN3 |
| PS50853 |  |  | 0.004262 | FN3 |
| IPR003961 |  |  | 0.000334 | FN3_dom |
| IPR009030 | 9.37E-05 | 0.005339 | 1.59E-05 | Growth_fac_rcpt_ |
| PS00856 | 0.02688 |  |  | GUANYLATE_KINASE_1 |
| PS50052 | 0.02688 |  |  | GUANYLATE_KINASE_2 |
| IPR013783 |  |  | 4.15E-05 | Ig-like_fold |
| IPR032695 |  |  | 0.0002 | Integrin_dom |
| IPR011009 | 0.04356 |  |  | Kinase-like_dom |
| IPR001791 |  |  | 8.36E-05 | Laminin_G |
| 2.60.40.10 |  |  | 4.19E-06 | NA* |
| 1.10.533.10 |  |  | 0.000225 | NA |
| PS51461 |  | 2.5E-05 |  | NC1_FIB |
| PF07714 | 0.04519 |  |  | Pkinase_Tyr |
| IPR000719 | 0.00585 |  |  | Prot_kinase_dom |
| PS00107 | 0.001612 |  | 0.00355 | PROTEIN_KINASE_ATP |
| IPR017441 | 0.01304 |  |  | Protein_kinase_ATP_BS |
| PS50011 | 0.006695 |  |  | PROTEIN_KINASE_DOM |
| PS00109 | 0.01171 |  |  | PROTEIN_KINASE_TYR |
| PS50001 |  |  | 0.005654 | SH2 |
| IPR008266 | 0.01171 |  |  | Tyr_kinase_AS |
| IPR020635 | 0.004333 |  |  | Tyr_kinase_cat_dom |
| SM00219 | 0.004333 |  |  | TyrKc |
| PS01208 | 0.04908 |  |  | VWFC_1 |

\*NA = Not available

**Table S4.** **Seed genes for the identification of a PDAC disease module with DADA**^13^. Genes are ordered following their genetic location; genes annotation shown as well.

| Gene | Chromosome | Description |
| --- | --- | --- |
| HDAC1 | 1 | histone deacetylase 1 |
| GLI2 | 2 | GLI family zinc finger 2 |
| TGFA | 2 | transforming growth factor alpha |
| IHH | 2 | Indian hedgehog signaling molecule |
| AREG | 4 | amphiregulin |
| HHIP | 4 | hedgehog interacting protein |
| EGF | 4 | epidermal growth factor |
| MAML3 | 4 | mastermind like transcriptional coactivator 3 |
| APC | 5 | APC regulator of WNT signaling pathway |
| HDAC2 | 6 | histone deacetylase 2 |
| CCND3 | 6 | cyclin D3 |
| EGFR | 7 | epidermal growth factor receptor |
| RAC1 | 7 | Rac family small GTPase 1 |
| SHH | 7 | sonic hedgehog signaling molecule |
| MYC | 8 | MYC proto-oncogene@ bHLH transcription factor |
| DKK4 | 8 | dickkopf WNT signaling pathway inhibitor 4 |
| DKK1 | 10 | dickkopf WNT signaling pathway inhibitor 1 |
| FRAT1 | 10 | FRAT regulator of WNT signaling pathway 1 |
| CCND1 | 11 | cyclin D1 |
| ATM | 11 | ATM serine/threonine kinase |
| MMP7 | 11 | matrix metalloproteinase 7 |
| FOSL1 | 11 | FOS like 1@ AP-1 transcription factor subunit |
| INHBE | 12 | inhibin subunit beta E |
| GLI1 | 12 | GLI family zinc finger 1 |
| WIF1 | 12 | WNT inhibitory factor 1 |
| IFNG | 12 | interferon gamma |
| ZIC2 | 13 | Zic family member 2 |
| JAG2 | 14 | jagged canonical Notch ligand 2 |
| SMAD3 | 15 | SMAD family member 3 |
| DLL4 | 15 | delta like canonical Notch ligand 4 |
| MAPK3 | 16 | mitogen-activated protein kinase 3 |
| SMAD4 | 18 | SMAD family member 4 |
| SMAD2 | 18 | SMAD family member 2 |
| SMAD7 | 18 | SMAD family member 7 |
| TGFB1 | 19 | transforming growth factor beta 1 |
| BMP2 | 20 | bone morphogenetic protein 2 |

**Table S5.** **Overlap between** individual perturbation expression profile (PEEP) for LT PDAC survivors with top 1% gene list from NetICS and DADA. NetICS= Network-based Integration of Multi-omics Data^25^; DADA= Degree-Aware Algorithms for Network Based Disease Gene Prioritization^13^.

| Identified in NetICS |  |  |
| --- | --- | --- |
| Patient ID | Nr* of PEEP genes in top 1% NetICS list | Top 5 genes |
| L1 | 1 | TNNI3 |
| L2 | 143 | TNNI3.IRS4.ACTN2.MAPK14.PLCG1. |
| L3 | 5 | FOSL1.MYL2.FCGR3A.USF2.NOSTRIN |
| L4 | 38 | ACTN2.PRKCA.ITGA8.AP2A1.NRAS |
| L5 | 20 | TJP2.ITGA10.SDC4.TYROBP.SPI1 |
| L6 | 26 | NPY.MPDZ.CNTN6.ITGB4.TNNC1 |
| L7 | 119 | TNNI3.HLA-DQA1.APOA4.IRS4.NPY. |
| L8 | 18 | GNAI3. ITGA10. KLRC3. SPI1.TNNC1 |
| L9 | 336 | ACTB. ACTG1.NFKB1.ITGB1.CTNNB1 |
| Identified in DADA |  |  |
| Patient ID | Nr of PEEP genes in top 1% DADA list | Top 5 genes |
| L2 | 8 | GLI2.RAC1.CUL1.ESR1.FBLN1.UBC.WNT11 |
| L3 | 1 | FOSL1 |
| L4 | 3 | TGFA.EGF.CDK2 |
| L5 | 2 | UBC.CDON |
| L7 | 9 | ESR1.GLI2.WNT11.GLI3.HDAC1.CDK2.UBC.CDON |
| L9 | 12 | GLI2.GLI3.UBC.RAC1.ACTB.AREG.DKK1.GRB2.MMP9.TGFB1.EGF.FOSL1 |

\*Nr=Number

**Table S6.**
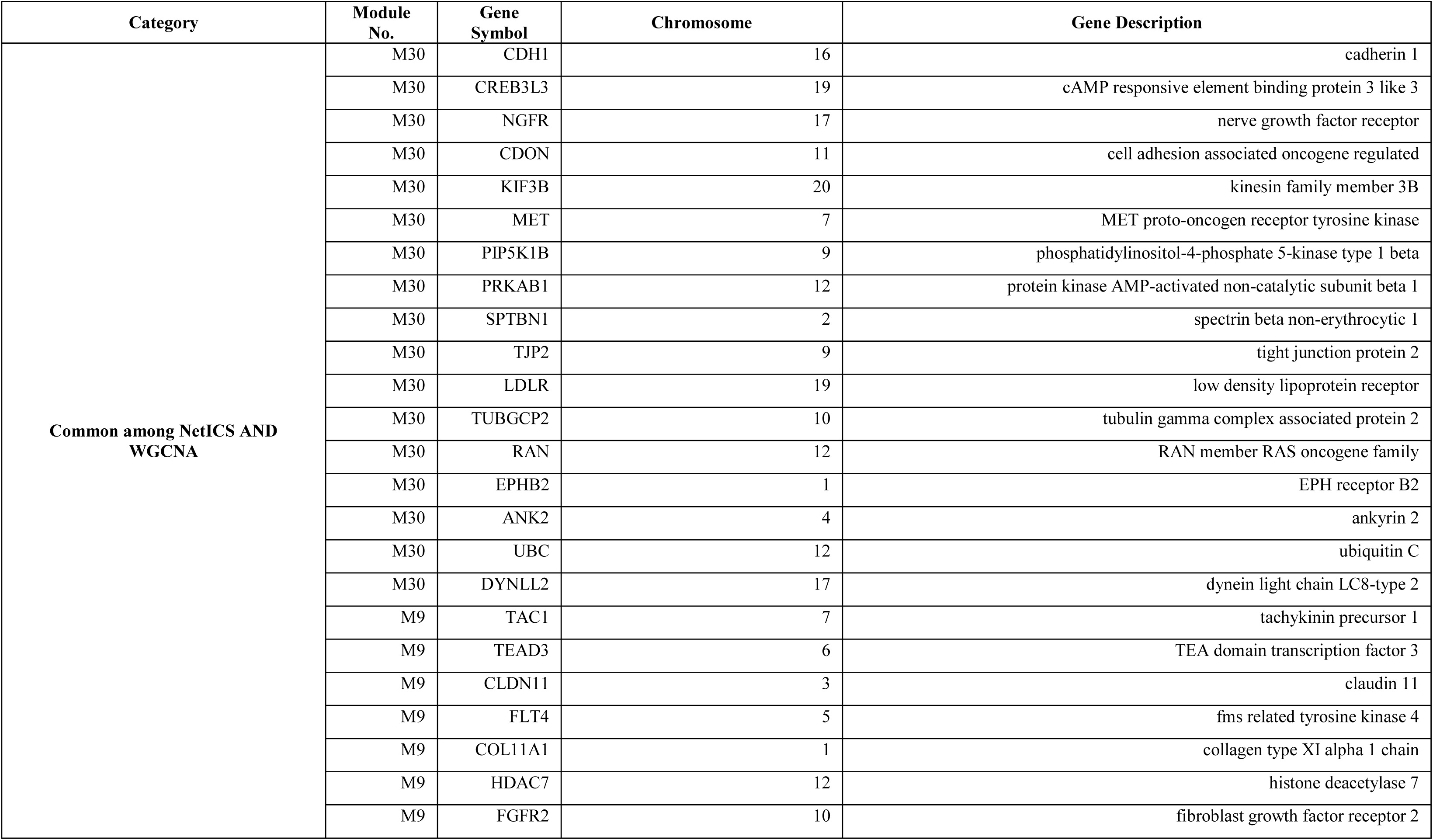
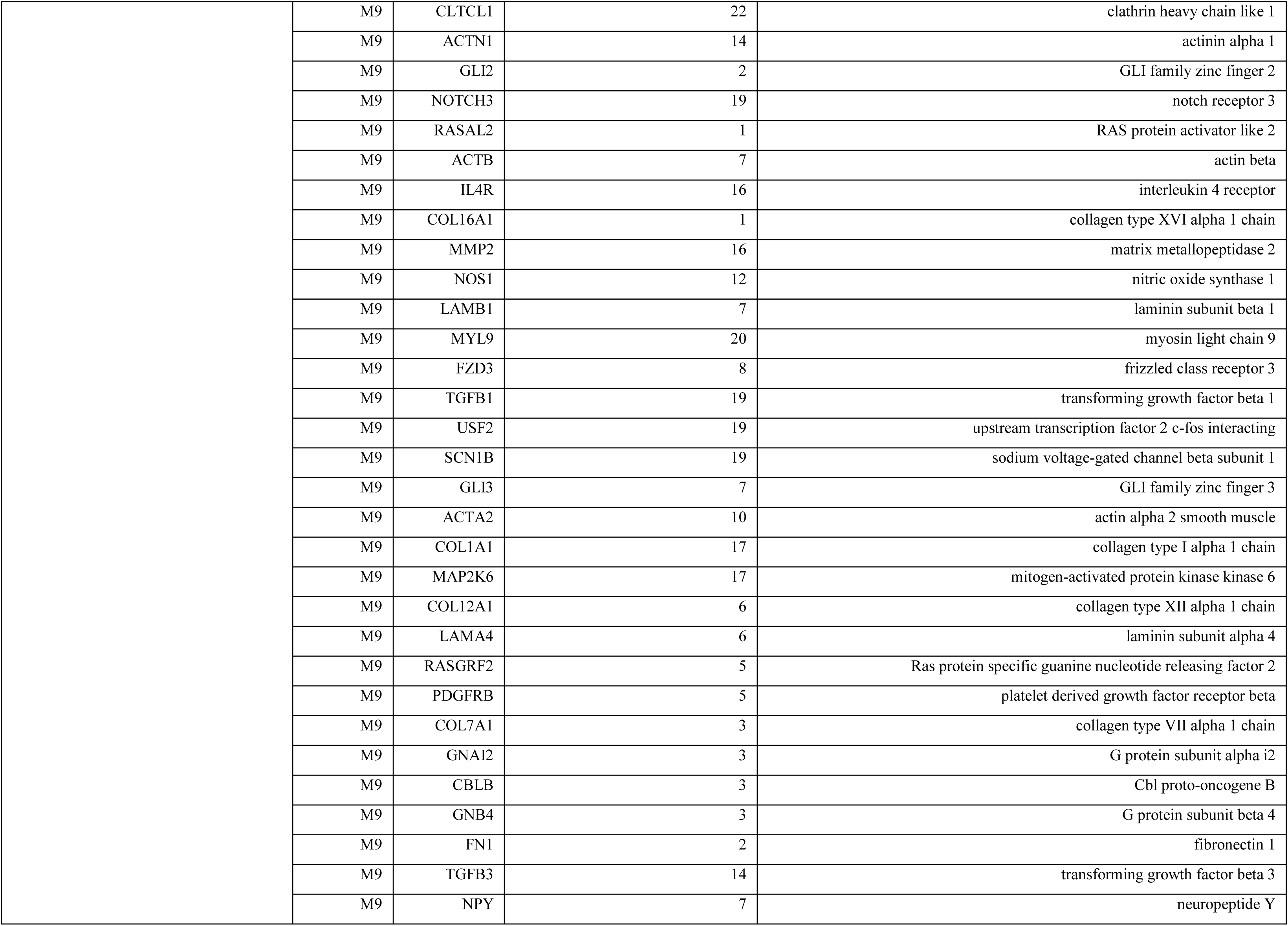
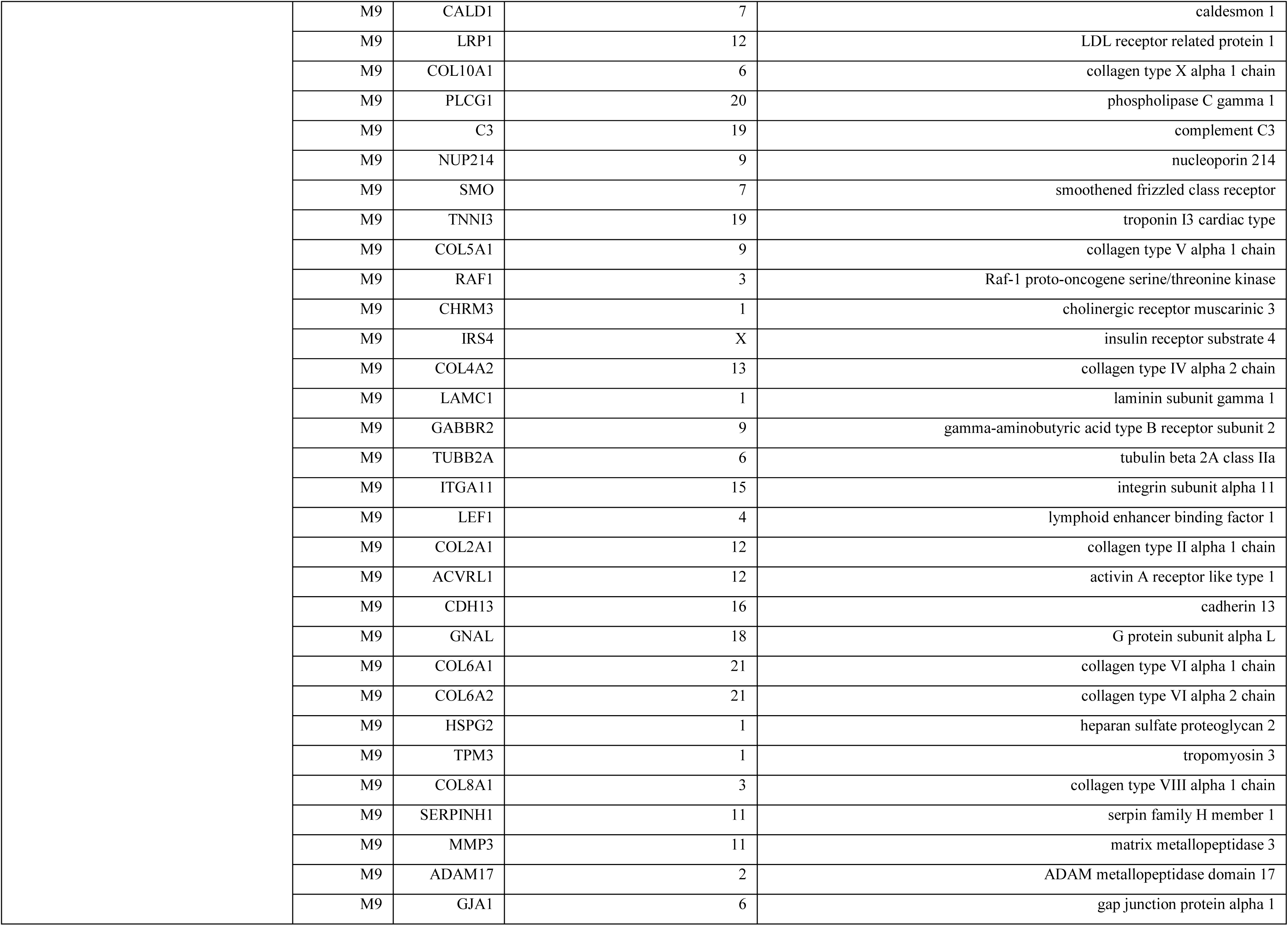
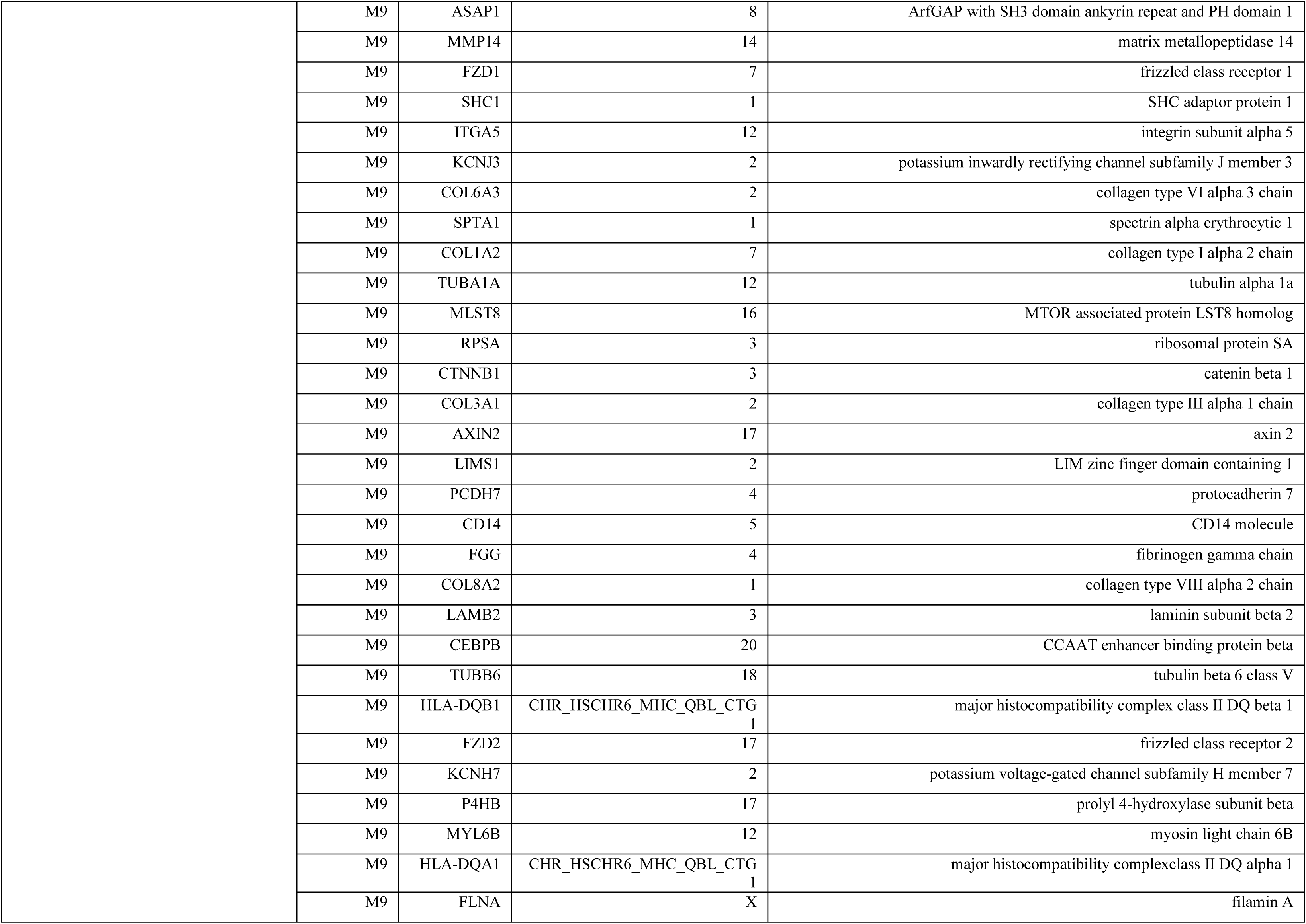
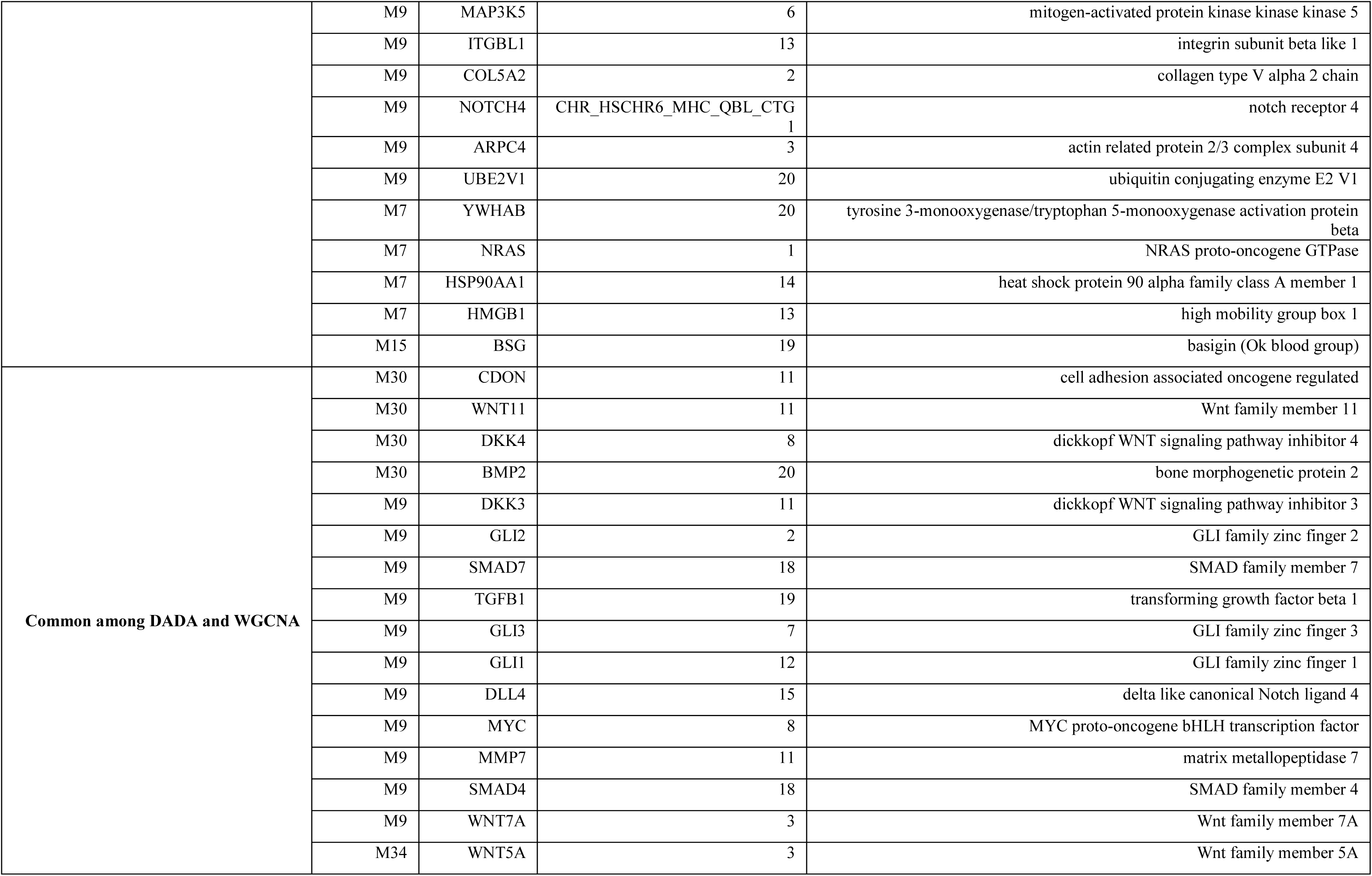
**Overlap between top 1% genes from NetICS /DADA and clinical modules, as defined in section 2.1 and 2.2.** WGCNA = Weighted correlation network analysis^10^; NetICS= Network-based Integration of Multi-omics Data^25^; DADA= Degree-Aware Algorithms for Disease Gene Prioritization^13^ .

**Figure S1.**
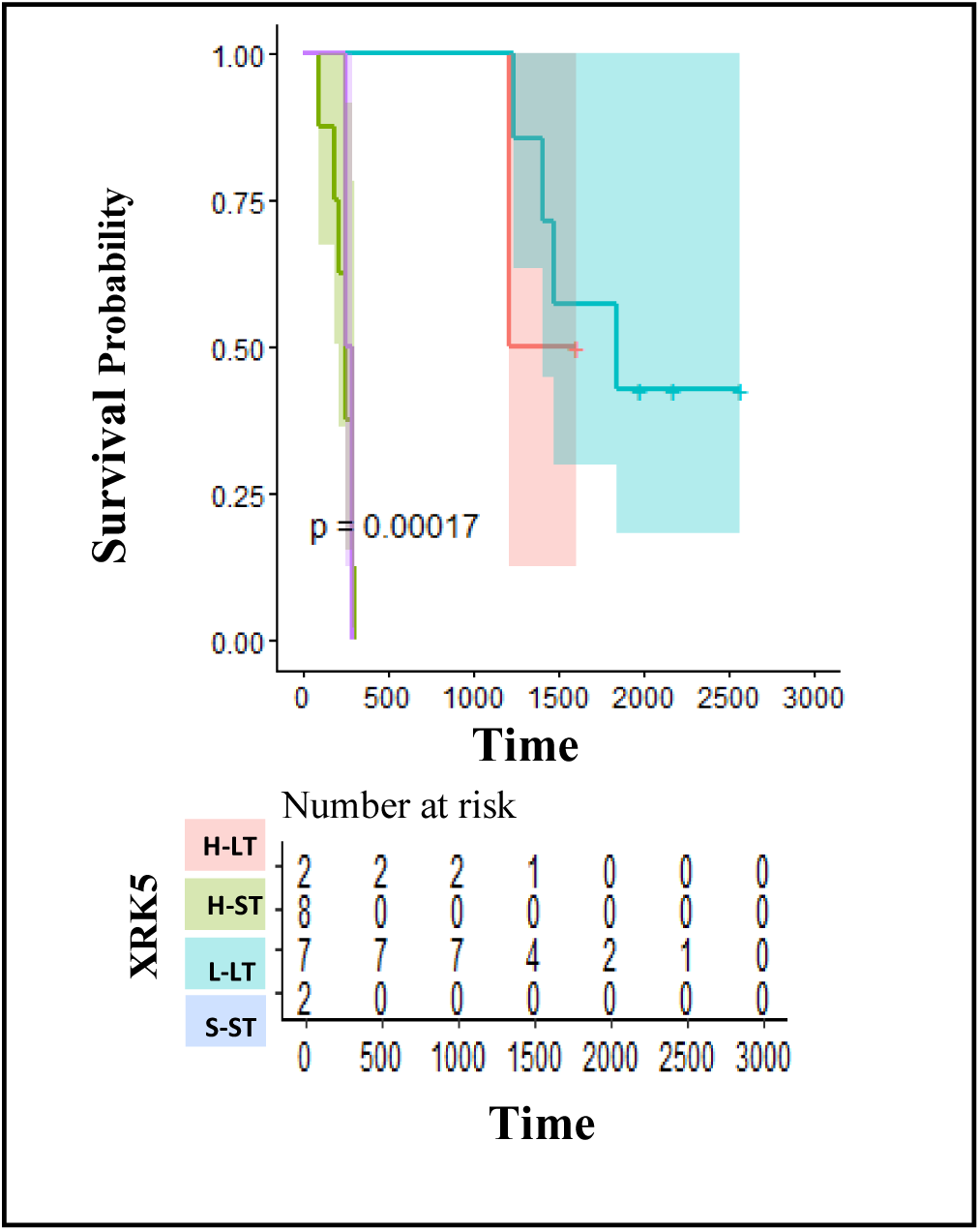
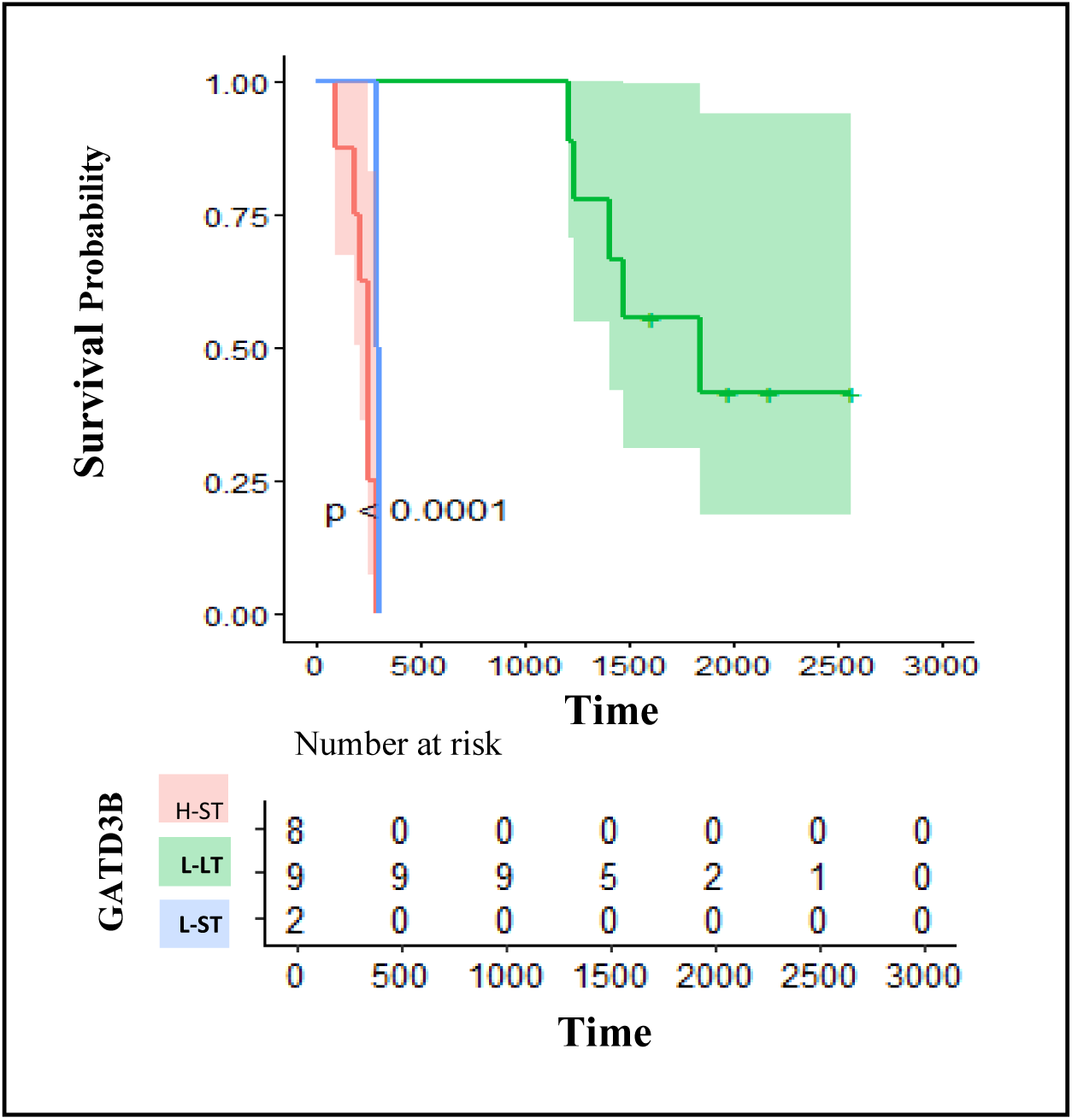
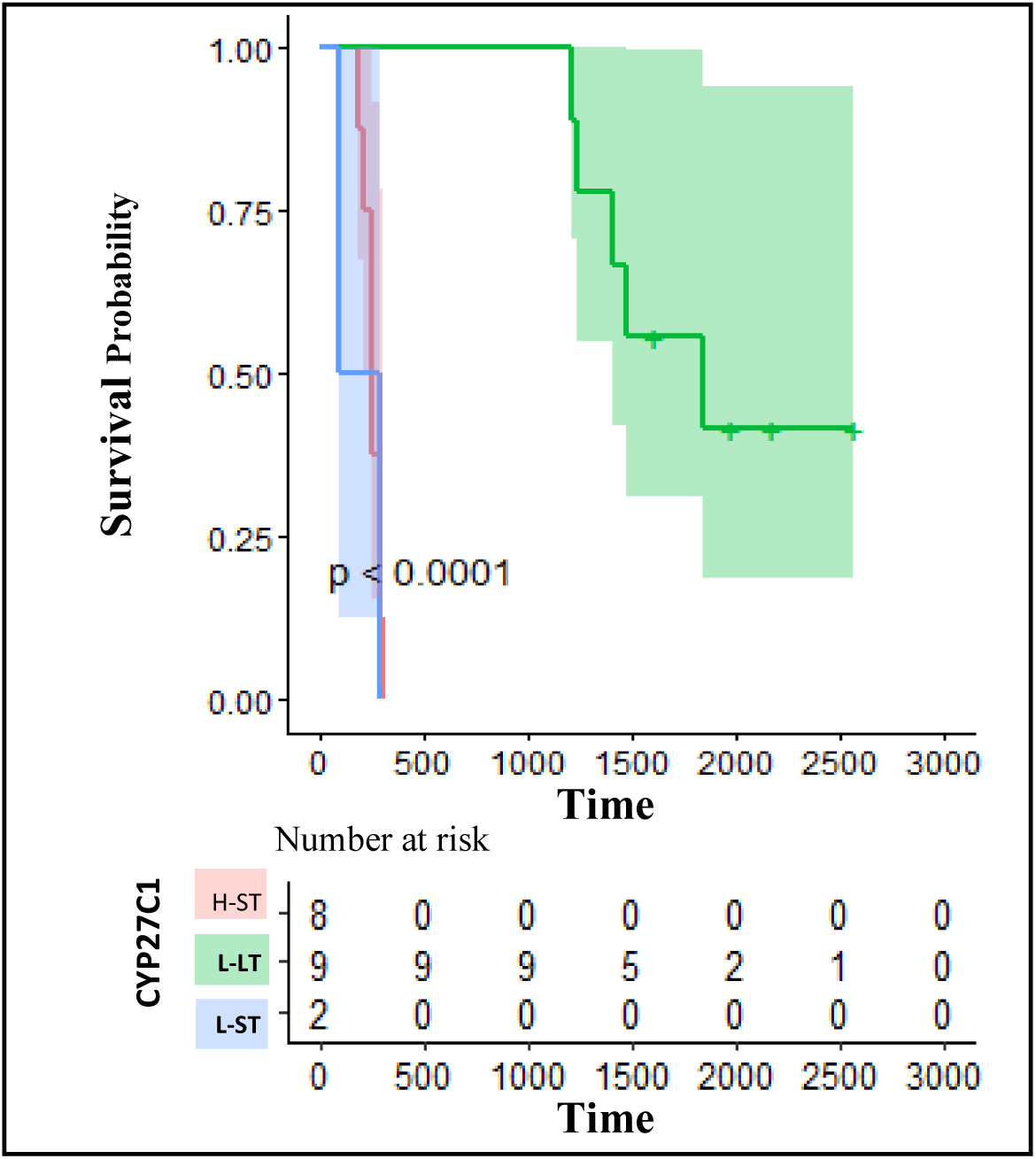
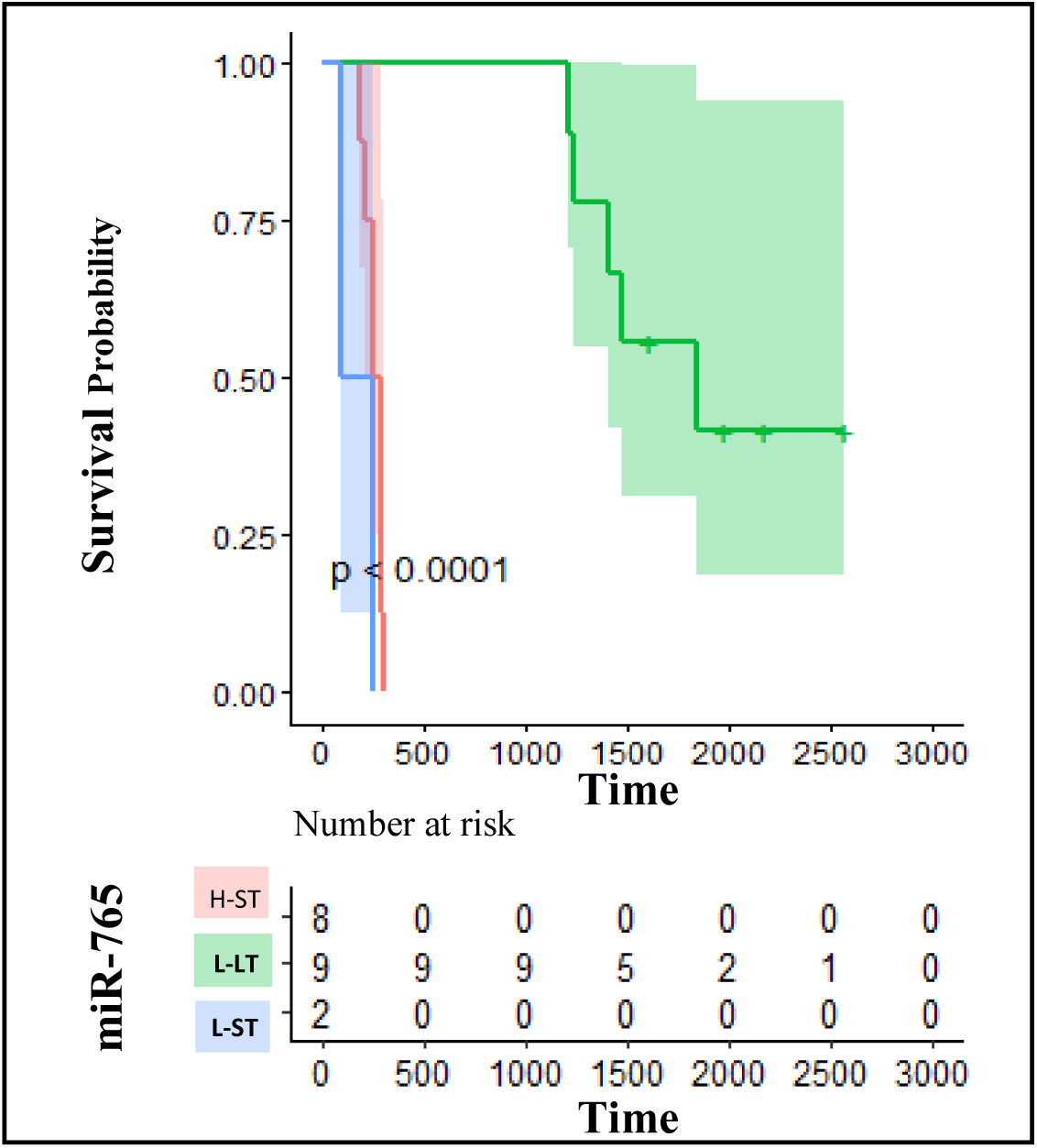
Kaplan-Meier plots for significant survival genes differentially expressed between long-term (LT: alive or death >=36 months) and short-term (ST: death >=3 months and <12 months) survival. A) Gene XRK5, p-value = 0.00017; B) Gene GATD3B, p-value < 0.0001; C) Gene CYP27C1, p-value < 0.0001) and D) Gene miR-765, p-value < 0.0001. Here, H and L represents the high and low expression groups identified (appendix 3).

**Figure S2.**
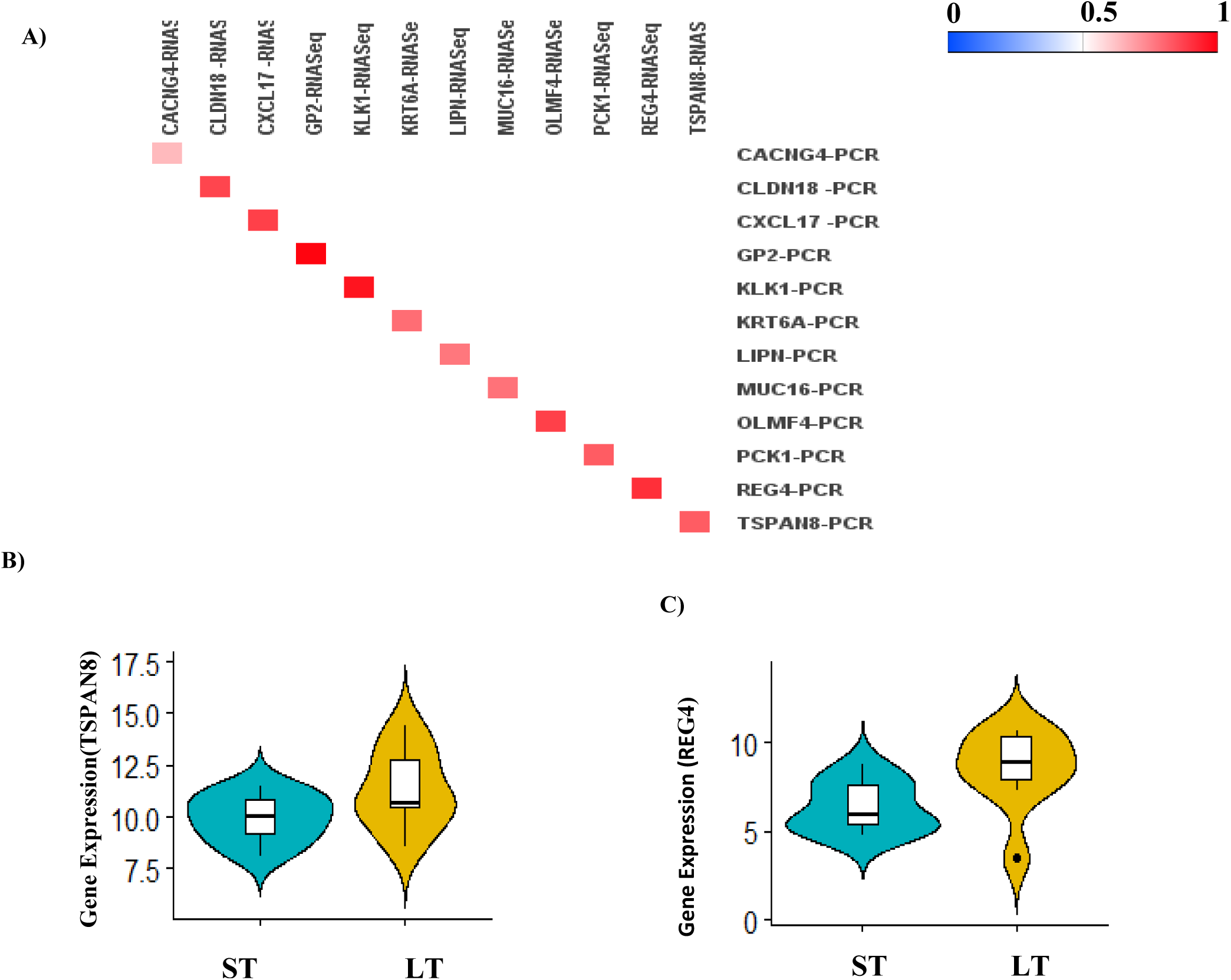
RNA-Seq supplemented with RT-PCR experiments: A) Correlation between RNA-Seq and RT-PCR expression values; B) Violin plot of TSPAN8 (gene ID: ENSG00000127324) by ST/LT PDAC survival; C) Same as B) but for REG4 (gene id: ENSG00000134193).

**Figure S3.**
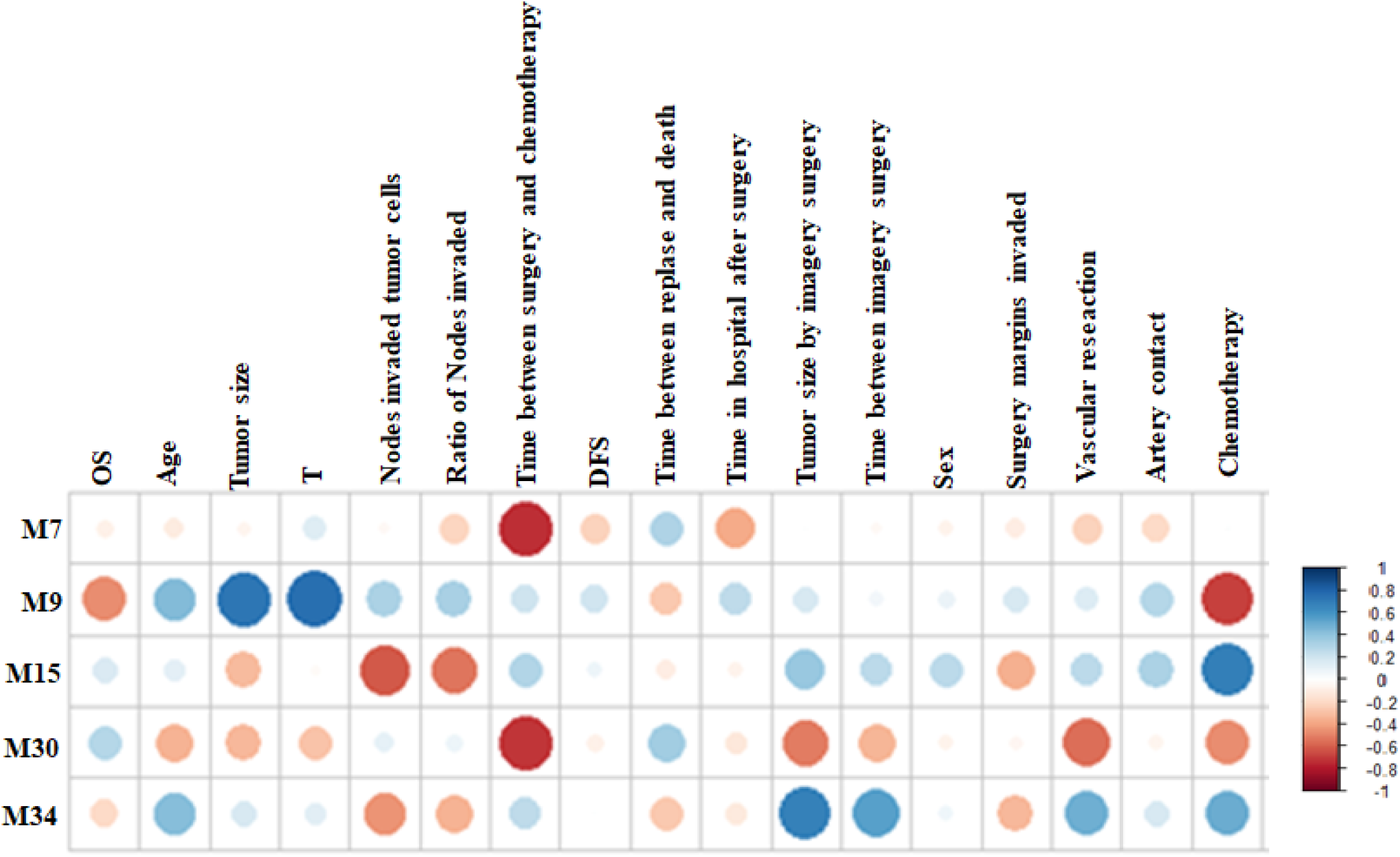
Clinical profile of WGCNA modules. Correlation patterns for WGCNA derived modules with significant association to clinical measurements (multiple testing adjusted p-value <0.05) are shown via corrplot^26^ (section 2.1). The more extreme the association (+1/-1) the deeper the color (dark blue/dark red). The sizes of the circles are proportional to the correlation coefficients. WGCNA = Co-Expression Network Analysis. Adjusted p values are not indicated in plot. Complete detail of significant modules is given in section: Survival group heterogeneity

**Figure S4.**
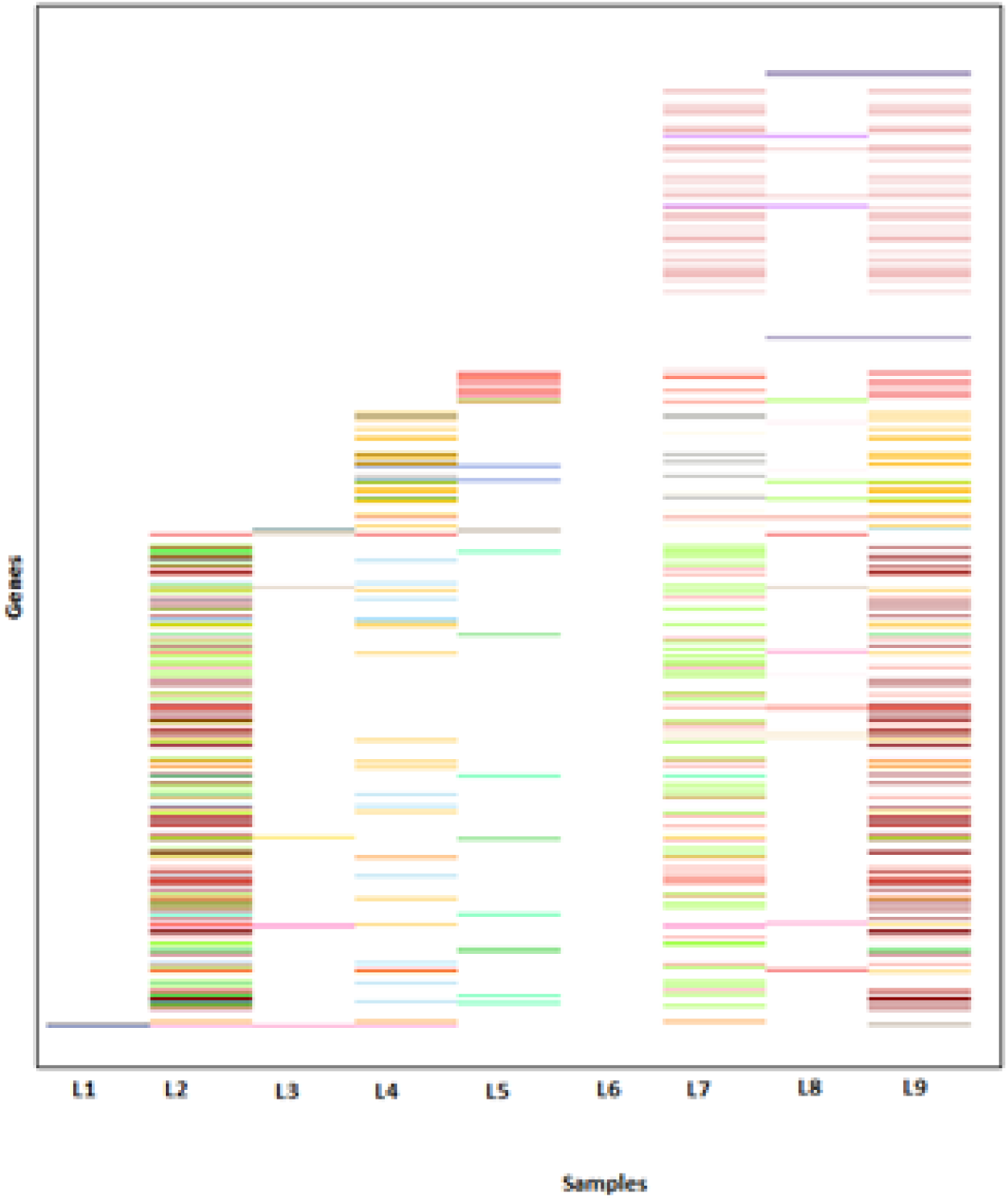
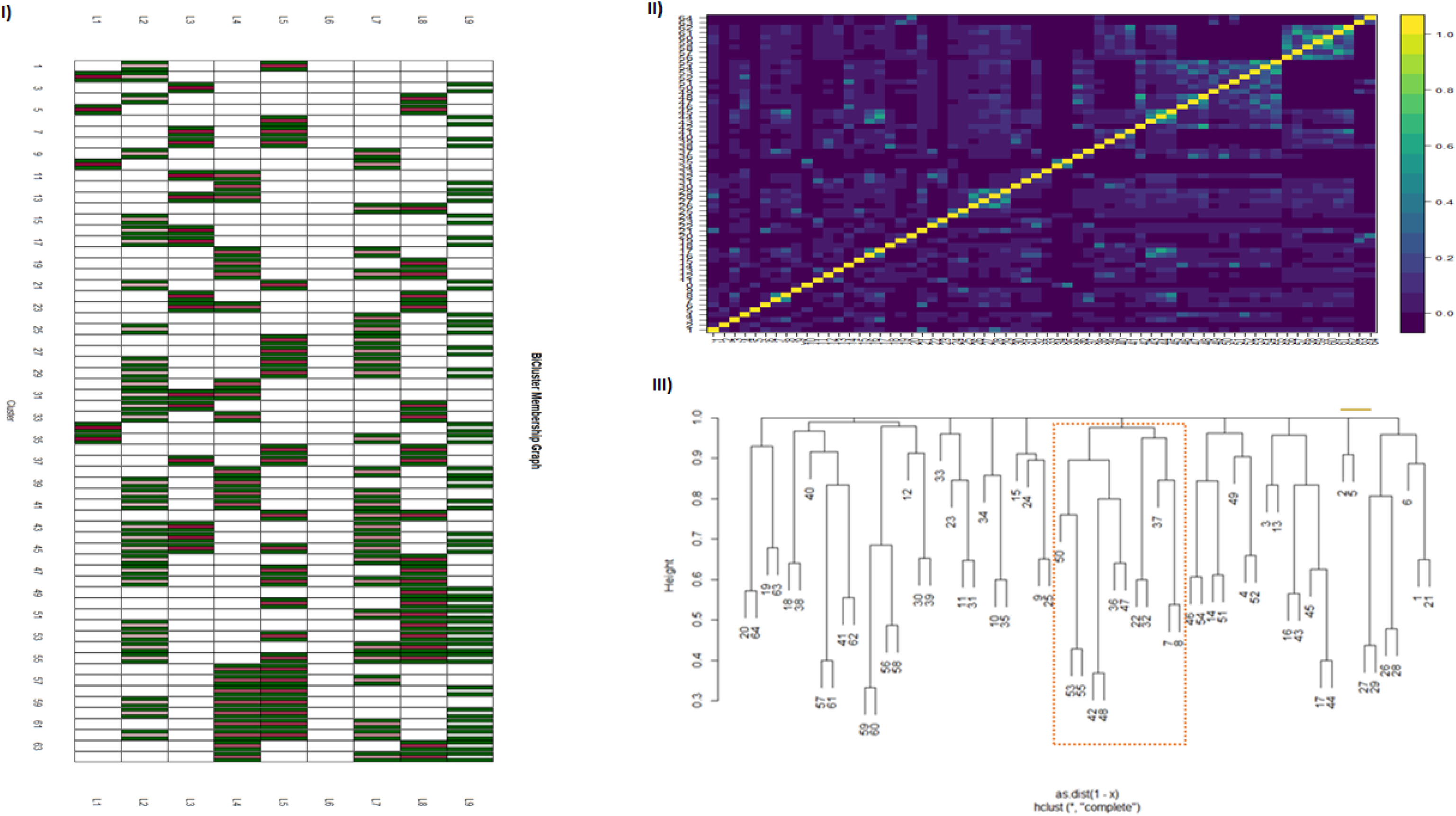
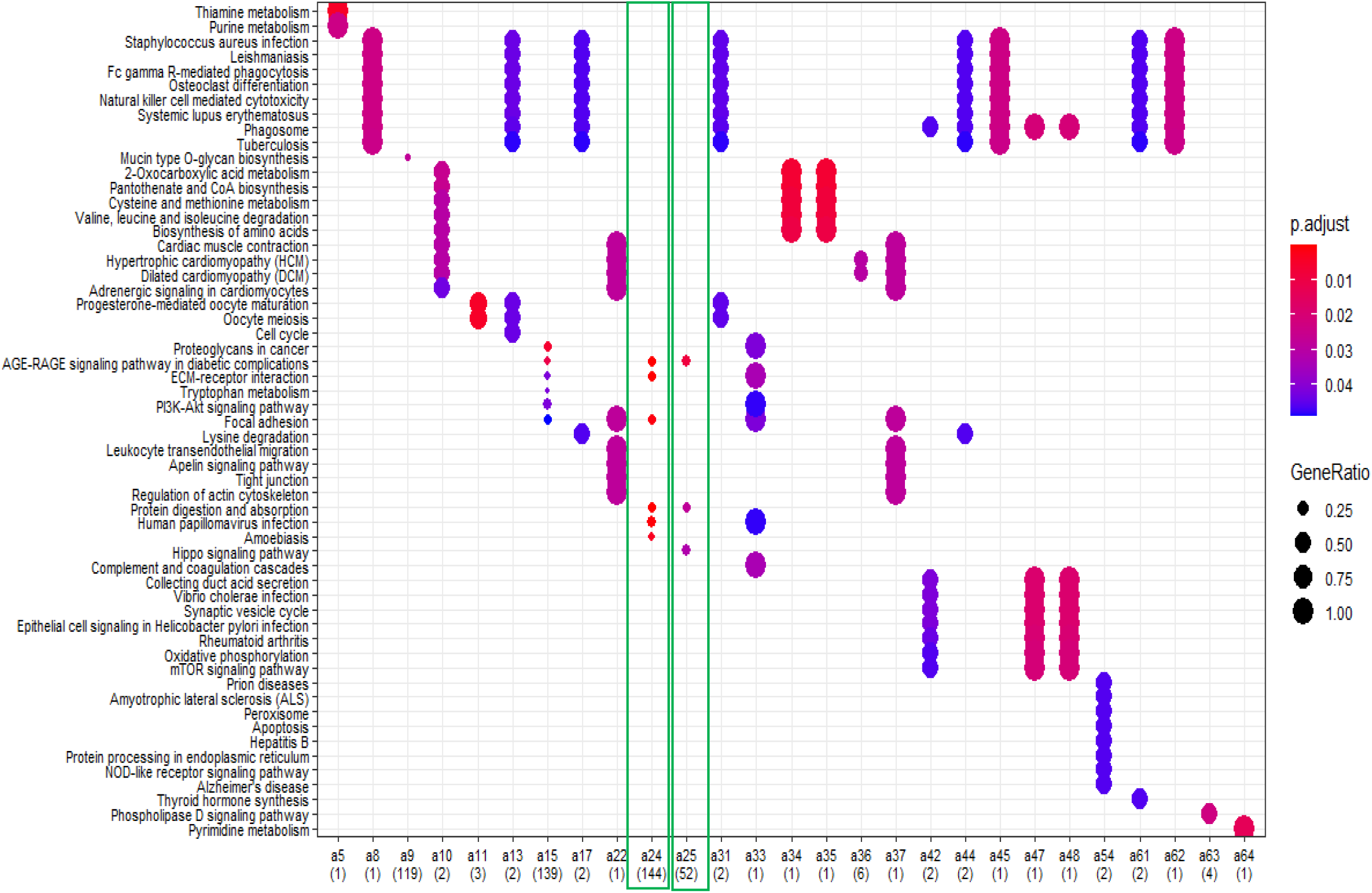
Bicluster analysis of perturbed genes in LT PDAC survivors: Input matrix is logical with genes (not) perturbed in an individual indicated by 1 (0). A) Sixty-four biclusters (BC) obtained from such a logical matrix (section 2.2) indicated as heatmap. Each color in heatmap represent cluster from 1 to 64. B) Advanced interpretation of identified biclusters via three different approaches. (I) **bicluster membership graph** based on BC cluster x LTS. (II) **Heatmap based on Jaccard similarity index** computed for the identified 64 biclusters ranging from 0 (no concordance) to 1 (perfect concordance). (III) **Hierarchical tree** constructed for the identified biclusters (appendix pp 4). C) Functional annotation of 64 biclusters with clusterProfiler^6^.

**Figure S5.**
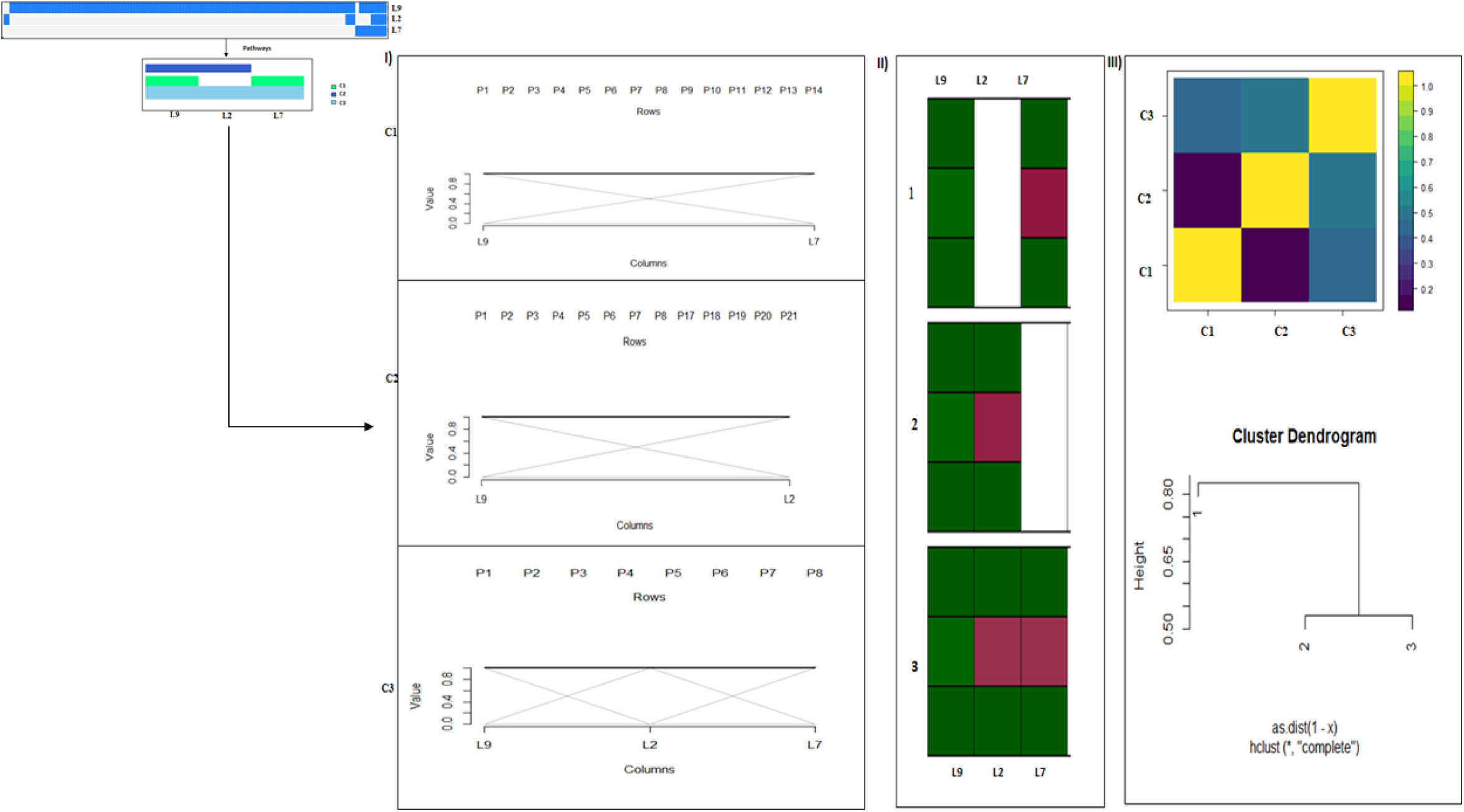
Bicluster analysis of enriched pathways in LT PDAC survivors: Input matrix is logical with enriched pathways (not) in an individual indicated by 1 (0). A) Three biclusters (BC) obtained from such a logical matrix (section 2.2). B) Advanced interpretation of identified biclusters via three different approaches. (I) **biclusters as lines of parallel coordinate graph** which indicates cluster specific detailed information. (II) **bicluster membership graph** based on BC cluster x LTS. (III) **Heatmap based on Jaccard similarity index** computed for the identified three biclusters ranging from 0 (no concordance) to 1 (perfect concordance). **Hierarchical tree** constructed for the identified biclusters (appendix pp 4).

**Figure S6.**
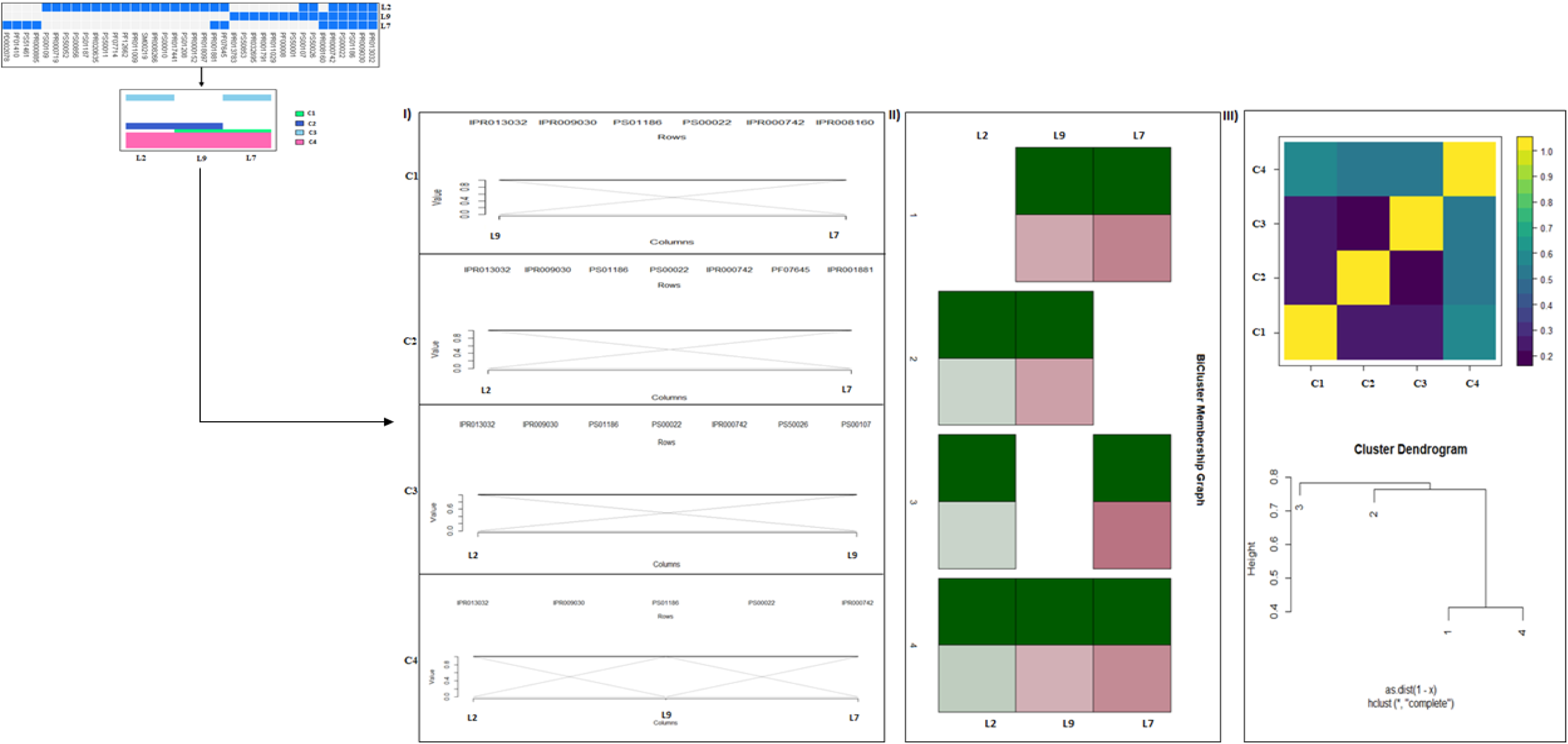
Bicluster analysis of enriched protein domains in LT PDAC survivors: Input matrix is logical with enriched protein domains (not) in an individual indicated by 1 (0). A) Four biclusters (BC) obtained from such a logical matrix (section 2.2). B) Advanced interpretation of identified biclusters via three different approaches. (I) **biclusters as lines of parallel coordinate graph** which indicates cluster specific detailed information. (II) **bicluster membership graph** based on BC cluster x LTS. (III) **Heatmap based on Jaccard similarity index** computed for the identified 4 biclusters ranging from 0 (no concordance) to 1 (perfect concordance). **Hierarchical tree** constructed for the identified biclusters (appendix pp 4).

**Figure S7.**
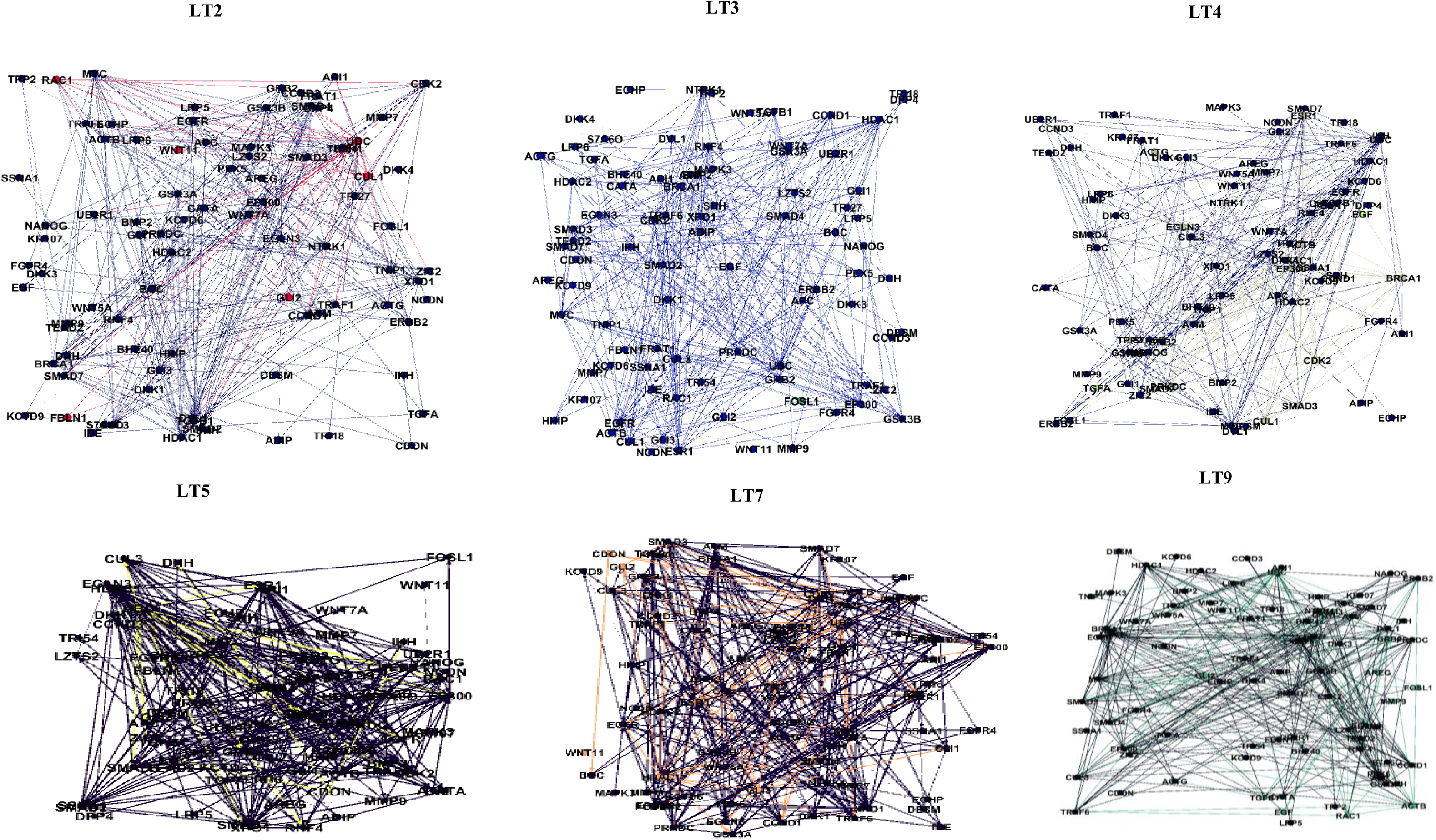
Superposition of PEEP induced perturbed genes to PDAC disease module derived via DADA. Individual-specific perturbed genes as identified by individual profiling with PEPPER^15^ (section 2.2) are highlighted with the same color per individual. LT2, LT3, LT4, LT5, LT7, and LT9 perturbed genes are indicated in red, green, light green, yellow, orange, and dark green, respectively. LT1, LT6 and LT8 specific perturbed genes showed no overlap in PDAC disease module derived via DADA (not shown in figure).

**Figure S8.**
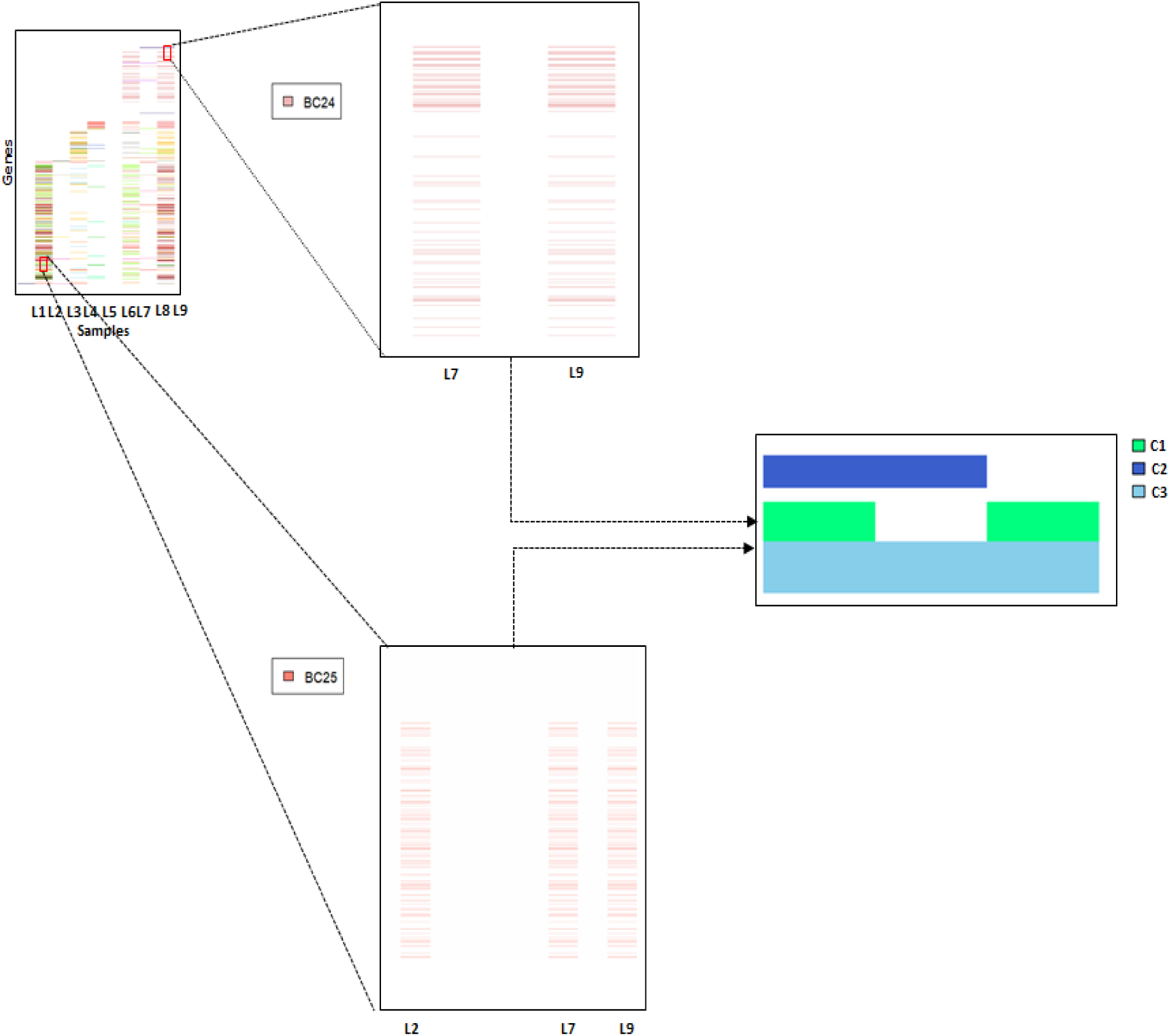
Aggregation of multi-level analyses. Genes present in groups BC24 and BC25 (Figure S4) were shown to be involved in pathway sets C2 and C3 (Figure S5) as indicated by the arrow (appendix 35).

## Notes

### Competing Interest Statement

The authors have declared no competing interest.

